# Membrane proteomics of the *Drosophila* circadian neural network

**DOI:** 10.64898/2026.05.20.726616

**Authors:** Chenghao Chen, Ratna Chaturvedi, Victoria Louis, Lauren E. North, Yongliang Xia, Maria Paz Gonzalez-Perez, Shaoxian Li, Vikas Kumar, Qing Yu, Patrick Emery

**Affiliations:** Department of Neurobiology, UMass Chan Medical School, 366 Plantation St, Worcester, MA 01605, MA; The Mass Spectrometry Facility, UMass Chan Medical School, 222 Maple Avenue, Shrewsbury, MA 01545; Department of Biochemistry and Molecular Biotechnology, UMass Chan Medical School, 364 Plantation St, Worcester, MA 01605, MA

## Abstract

The *Drosophila* brain contains approximately 240 pacemaker neurons that control circadian behaviors. Their molecular clocks are synchronized, but the phase of their rhythmic activity differs, so that specific groups of clock neurons can properly time various behaviors. To understand in depth how the circadian neural network functions, we surveyed its membrane proteome in the morning and evening, using *in situ* protein labeling coupled with mass spectrometry. We detected known regulators of circadian behavior and identified novel membrane or membrane-associated proteins in clock neurons. Through genetic screens, we found that many of these proteins regulate circadian behavior. In particular, Octβ1R modulates evening locomotion under long photoperiod, while Piezo regulates morning activity under short photoperiod. Loss of Piezo alters rhythmic neuronal activity and compromises the structural plasticity of the neurons controlling locomotion at dawn. Our work thus illustrates the power of proteome-guided genetic screening to understand the mechanisms underlying circadian behavior.

## Introduction

Circadian rhythms are a nearly universal feature of life, allowing organisms to optimize their metabolism, physiology, and behavior in alignment with the solar day^1^. They are thus critical for the health and survival of most organisms. In humans, the chronic misalignment of these internal rhythms with environmental cycles is an important contributor to various metabolic and psychiatric disorders^2–4^. Circadian behavior is controlled by specific brain neurons. In mammals, it is driven by pacemaker neurons of the suprachiasmatic nucleus (SCN)^5^. In *Drosophila melanogaster*, circadian locomotor behavior is governed by a functionally and molecularly homologous network of approximately 240 pacemaker neuron^6^. Within these cells, an evolutionarily conserved molecular clock operates through a series of transcriptional feedback loops^7^. In brief, the transcriptional activators Clock (CLK) and Cycle (CYC) drive the expression of *period* (*per*) and *timeless* (*tim*); subsequently, PER and TIM proteins translocate to the nucleus to repress CLK/CYC activity. Interestingly, there is a division of labor among *Drosophila* circadian neurons. For example, the small ventral lateral neurons (s-LNvs) control circadian behavior in constant darkness (DD) and also drive circadian locomotor activity in the morning under a light:dark (LD) cycle^8–11^. They are therefore also referred to as Morning (M-) oscillators. On the other hand, the dorsal lateral neurons (LNds) and an additional s-LNv promote evening activity and are thus called Evening (E-) oscillators ^10,11^. Dorsal neurons (DN1-3) exert various functions, such as modulating rhythmic behavior as a function of light and temperature conditions and regulating rhythmically sleep or temperature preference^12–15^.

The molecular circadian clock oscillates with the same phase in almost all subsets of circadian neurons: PER and TIM levels are low at the end of the day, and high late at night. Nevertheless, these neurons exhibit distinct phases of electrical activity^16^ and Ca^2+^ oscillations^17^, which presumably explain how they control specific behaviors. For instance, the M-oscillators are preferentially active at dawn to drive morning locomotor activity, whereas the E-oscillators fire most robustly at dusk. Although Ca^2+^ rhythms are known to be dampened or abolished in molecular clock mutants^17^, the link between the cytoplasmic/nuclear molecular oscillator and desynchronized neural activity is still poorly understood. The principal mechanism that has been proposed is called the “bicycle model”, which appears to apply to mammalian SCN neurons as well^18^. In *Drosophila*, the Na^+^ leak channel *narrow abdomen* (NA) promotes depolarization of DN1p neurons and l-LNvs in the morning, while K^+^ channels such as Shaw and Shal modulate currents at night in the l-LNvs^18,19^. These opposing oscillations in Na^+^ and K^+^ currents contribute to daily rhythms in neuronal activity. The bicycle model is, however, insufficient to explain the wide range of activity rhythms observed in clock neurons. For example, while the l-LNvs and the DN1ps share NA as a rhythmic regulator of activity, their activity peaks are not synchronized. In addition to cell-autonomous rhythms impacting their electrophysiological properties, circadian neuron activity is modulated by neuronal communications within the circadian neural network^20^, and by input pathways^21,22^. Thus, we reasoned that the orchestration of circadian firing patterns is mediated by the cell-membrane proteome and its dynamic daily remodeling.

Cell-membrane proteins (CMPs)—including ion channels, neurotransmitter and neuropeptide receptors, and adhesion molecules—are the primary effectors of neuronal communication and excitability. However, profiling the CMP landscape with high spatiotemporal resolution within intact tissues remains a significant technical challenge, particularly in *Drosophila*, given the small number of neurons involved, and their minute size. Transcriptomic approaches, particularly at the single-cell level, have proven effective at defining the molecular characteristics of circadian neurons, including CMPs^23–25^. However, transcriptional profiles do not necessarily correlate well with protein expression, particularly when examining specific subcellular compartments^26–28^. Consequently, a direct cell-type-specific, *in situ* characterization of the clock neuron membrane proteome is essential to understand comprehensively how circadian neural activity and ultimately rhythmic behavior are generated.

Here, we employed a combination of cell-type-specific proximity biotin labeling and quantitative tandem mass tag (TMT) proteomics to snapshot the cell-membrane proteome of *Drosophila* clock neurons at dawn and dusk. Our analysis revealed a predominant enrichment of specific CMPs at dawn, coinciding with daily peaks in structural plasticity and neural activity. Through a proteome-informed *in vivo* genetic screen, we identified several novel regulators of circadian rhythms, including components of the exocytosis machinery that maintain circadian rhythmicity, and ion channels that regulate period length. Furthermore, we found that the G-protein coupled receptor (GPCR) Octβ1R regulates the evening activity peak under long photoperiod, while the mechanosensitive channel Piezo^29,30^ modulates daily clock neuron activity, structural plasticity of M-oscillators and behavioral adaptation to short photoperiods. Together, these data provide the first comprehensive cell-membrane proteomic landscape of circadian pacemaker neurons, offering a molecular roadmap for understanding how the circadian clock translates temporal information into rhythmic behavior.

## Results

### Cell-type-specific membrane proximity labeling platform in clock neurons

To understand in depth how the *Drosophila* circadian neural network controls behavior, we aimed to probe its membrane proteome. However, characterizing the membrane proteome of specific brain neurons is challenging due to the dense intermingling of neurites, synaptic connections and glial processes. This difficulty is compounded in *Drosophila* by the minute size of its brain and neurons. The neuron-type specific expression of an extracellular Horseradish Peroxidase (HRP) fused to CD2 for membrane tethering has been successful in identifying relevant membrane proteins in the *Drosophila* antennal lobe^26^. However, a caveat with this method is the labeling of proteins on the membranes of cells that are adjacent to the target neurons^31^. To achieve high-fidelity, cell-autonomous labeling, we decided to utilize a second-generation ascorbate peroxidase (APEX2) fused to the intracellular C-terminus of the rat transmembrane protein CD8^32^ (**Figure 1A**). APEX2 offers significant advantages over biotin ligase-based methods (e.g., BioID), including faster reaction kinetics (<1min) and a smaller labeling radius, which provides a high-resolution temporal snapshot of the membrane proteome^33^. Importantly, APEX2’s intracellular location should enable the identification of not only neuronal transmembrane proteins, but also proteins localized close to the membrane, within the APEX2 labeling radius. Those could include important modulators of membrane proteins such as cytoskeletal proteins, or factors implicated in signal transduction, for example, downstream of neurotransmitter receptors. To study the membrane proteome of clock neurons, we thus decided to express CD8-APEX2 with the *Clk-856-GAL4* driver (here abbreviated to *Clk-GAL4*), which is expressed in most circadian neurons^25,34^. The workflow would be to 1) dissect and expose brains to H_2_O_2_ and biotin-phenol (BP) to induce *in situ* biotinylation, 2) lyse the brains and incubate the lysates with streptavidin beads to enrich the labeled proteins, 3) trypsin-digest the purified proteins and TMT-label the resulting peptides, and 4) use liquid chromatography-tandem mass spectrometry analysis (LC-MS/MS) for peptide identification and quantification **(Figure 1A)**.

**Figure 1.**
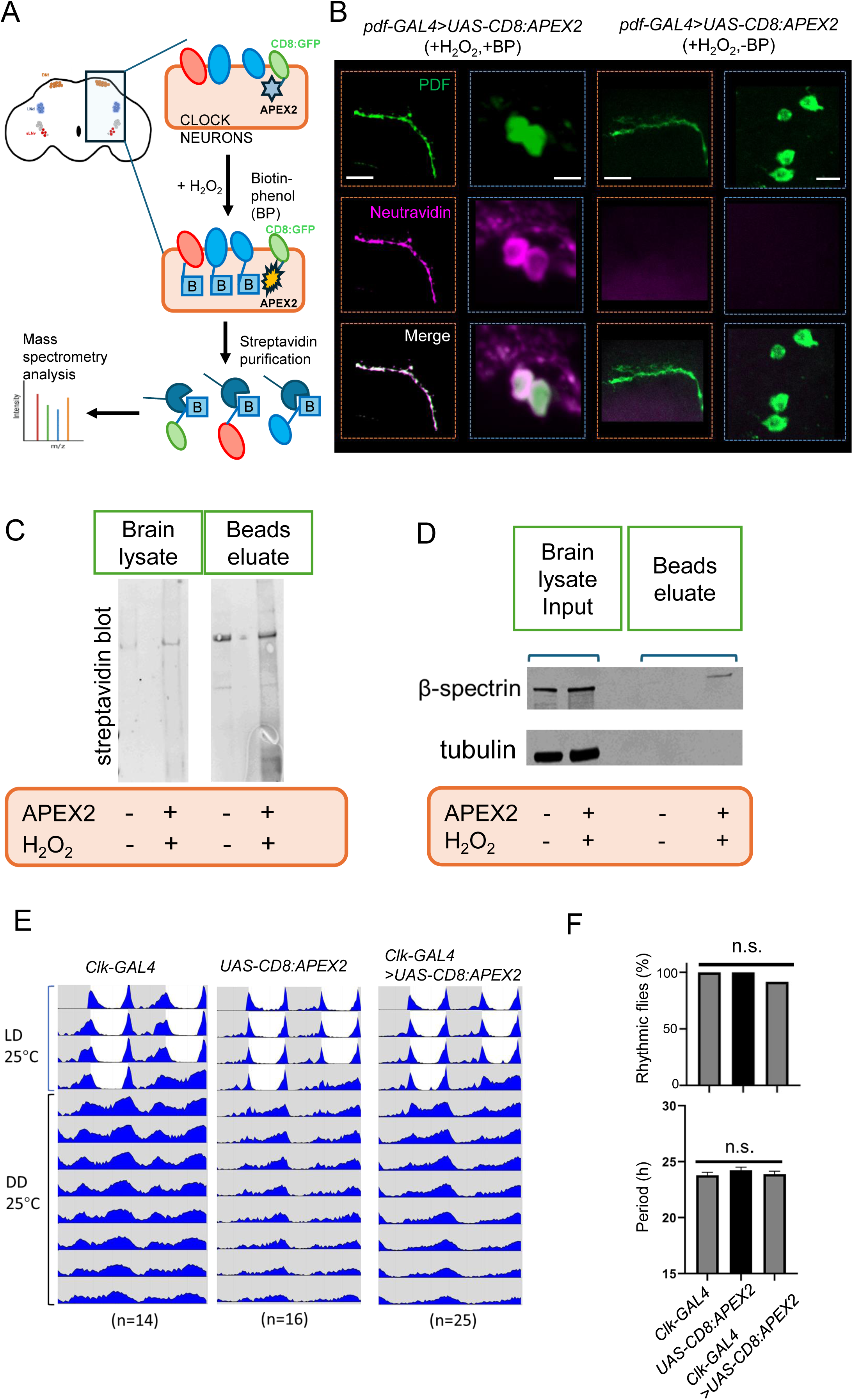
Cell-membrane proximity labeling of clock neurons in intact *Drosophila* brains. (A) Schematic of proximity biotinylation and mass spectrometry-based proteomic profiling in intact brains. In the presence of H_2_O_2_ and biotin-phenol (BP), a plasma membrane-tethered, intracellular-facing ascorbate peroxidase (APEX2) catalyzes the biotinylation of nearby membrane protein (red and blue). Labelled proteins are extracted and enriched using Streptavidin beads followed by quantitative mass-spectrometry analysis. (B) Representative confocal images of experimental (+BP, +H_2_O_2_) and negative control (-BP) conditions. Neutravidin (magenta) and PDF (green) staining reveals robust biotinylation signals at the s-LNv axonal projections (left) and somata (middle) only under experimental conditions. Scale bars, 10 μm. (C) Streptavidin blots of whole-brain lysates (left) and post-enrichment bead eluates (right). Note the marked enrichment of biotinylated proteins in experimental samples expressing APEX2 in most circadian neurons with *Clk-GAL4* (+APEX2, +H_2_O_2_) compared with controls (-APEX2, - H_2_O_2_). (D) Immunoblots for the cytoplasmic marker tubulin and the cytoskeletal and membrane-associated protein β-spectrin in whole-brain lysates and post-enrichment eluates. (E) Targeted APEX2 expression in clock neurons does not impair circadian behavior. Representative double-plotted actograms of flies with the indicated genotypes are shown. Flies were entrained (LD, 25°C) for 4 days before release into constant darkness (DD, 25°C) for 8 days. (F) Quantification of the percentage of rhythmic flies and period length. N are indicated in brackets; n.s., not significant by a Fisher exact test (percentage rhythmicity) and a Kruskal–Wallis test followed by Dunn’s multiple-comparisons test (period).

We first verified the specificity of our labeling approach. We expressed *CD8-APEX2* only in a subset of circadian neurons, the ventral lateral neurons (LNvs), using *pdf-GAL4*^8^. After dissection, the brains were exposed to H_2_O_2_ and BP to initiate the proximity-labeling reaction, and fluorophore-conjugated neutravidin was used to visualize APEX2-mediated biotinylation. As predicted, the fluorescent signal was restricted to the LNvs, and could clearly be detected on both their somata and neuronal projections (**Figure 1B**, left). Importantly, APEX2 expression and the labeling reaction did not induce gross morphological defects, as confirmed with an antibody to PDF, a neuropeptide specific to the LNvs^8^. Control experiments omitting BP showed no detectable biotinylation (**Figure 1B**, right).

We then expressed CD8-APEX2 with *Clk-GAL4*, dissected a small number of brains (ca. 100) and triggered biotinylation before extracting proteins. Probing of whole-brain lysates purified on streptavidin beads by Western blots confirmed the robust enrichment of biotinylated proteins in APEX2-expressing samples compared with controls (**Figure 1C**). Immunoblotting further indicated successful membrane-associated protein enrichment. β-spectrin, a cytoskeletal protein localized close to the plasma membrane, was enriched in the eluate, whereas the cytosolic protein tubulin was not detectable (**Figure 1D**). Importantly, flies expressing CD8-APEX2 in most clock neurons exhibited normal locomotor rhythms in constant darkness (DD), indicating that expression of this fusion protein does not disrupt circadian neuron function (**Figures 1E–F**). In summary, these important control experiments demonstrated the specificity of the biotin labeling to clock neurons, and the feasibility of enriching and identifying their membrane and membrane-associated proteins.

### TMT-based quantitative proteomics reveals temporal shifts in the clock proteome

We then performed time-dependent proteomic profiling using *Clk-GAL4*. We dissected brains at two distinct time points: ZT1 (dawn), when the majority of pacemaker neurons are active, and ZT11 (dusk), a period of relative quiescence for most clock neurons except the E-oscillators (LNds)^17^. To better quantify protein changes and filter out contaminants captured in negative controls, we employed a tandem mass tag (TMT) strategy^35^, incorporating two biological replicates and their negative controls (without APEX2 expression and H_2_O_2_ in the reaction) per time point, and an additional channel served as a “pool control” (**Figures 2A and 2C**). For each sample, about 800 brains were dissected and treated with BP. Their biotinylated proteins were purified and analyzed by TMT-based quantitative mass spectrometry.

**Figure 2.**
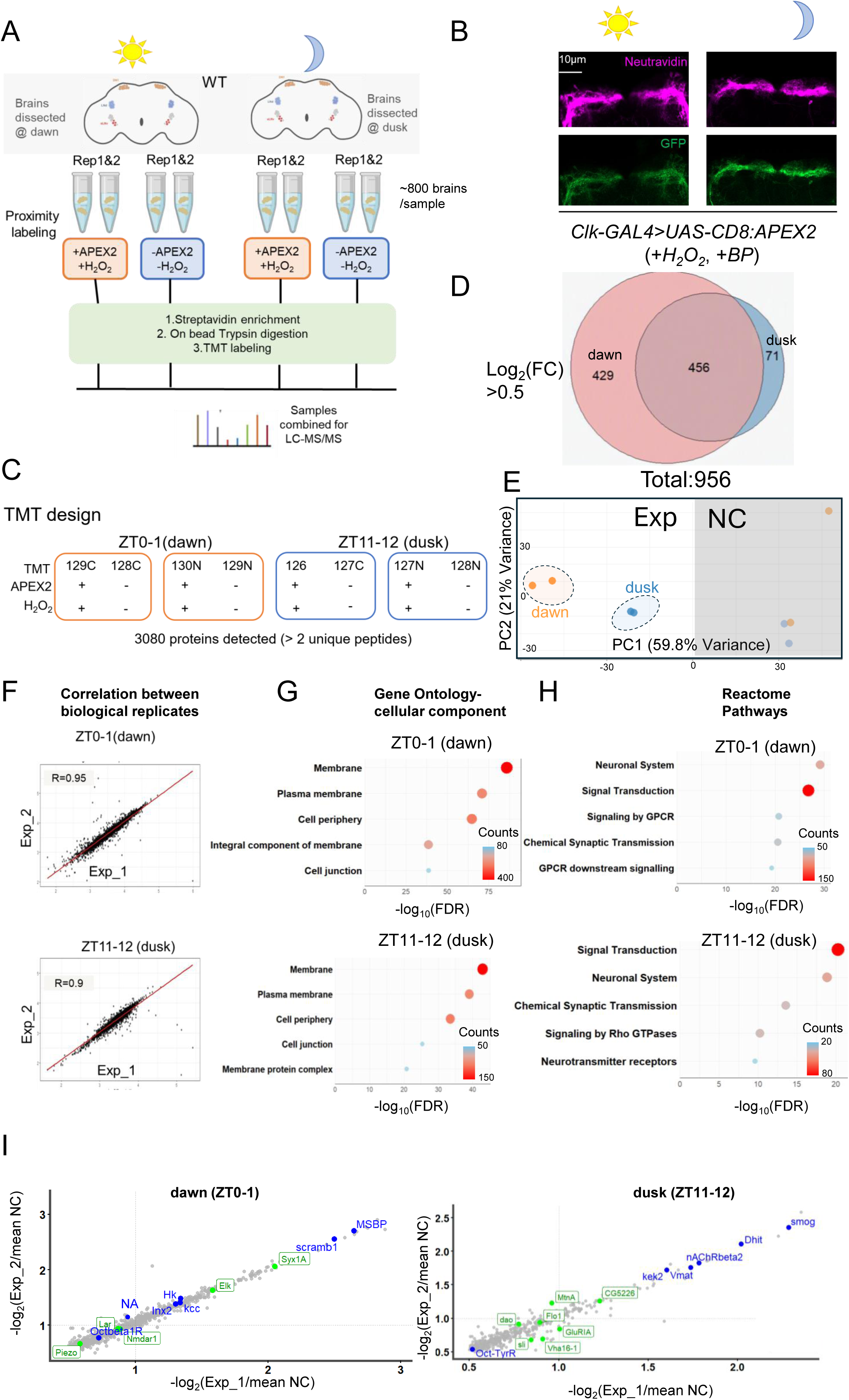
Quantitative TMT-based proteomics reveals time-dependent shifts in the clock neuron surface proteome. (A) Experimental workflow for time-dependent membrane proteomic profiling in clock neurons. (B) Representative confocal images of experimental flies at two time points. Neutravidin (magenta) and GFP (green) staining show that the total levels of biotinylated surface proteins do not obviously oscillate across time points. Scale bars, 10 μm. (C) Design of the tandem mass tag (TMT)-based quantitative proteomic experiment, pairing each experimental sample with a negative control. (D) Venn diagram of the clock neuron proteomes after ratiometric analyses. The dataset comprises 956 proteins, with 456 shared across both time points. (E) Principal-component analysis (PCA) of TMT reveals distinct, time-dependent clustering of experimental samples. Exp: experimental group: dawn (129C and 130N) and dusk (126 and 127N), NC: negative control group: dawn (128C and 129N) and dusk (127C and 128N) (F) Correlation analysis of biological replicates at dawn and dusk. (G) Gene Ontology (GO) analysis showing enrichment in cellular compartment terms associated with the plasma membrane. (H) Top 5 terms from Reactome analysis, highlighting events relevant to membrane protein function. (I) Enrichment of proteins in clock neurons at dawn (ZT0-1) and dusk (ZT11-12). Blue, examples of membrane proteins found at both time points; green, examples of unique membrane proteins enriched only at a single time point. See also **Data S1**.

A total of 3080 proteins were identified with >2 unique peptides after mass-spectrometry analysis (**Figure 2C; Data S1**). To remove endogenous biotinylated proteins as well as non-specific binding to Streptavidin beads, we applied a stringent ratiometric cutoff, with the TMT ratio between experimental and control samples having to be greater than 1.41(log_2_FC>0.5). This yielded 885 membrane-associated proteins enriched at dawn and 527 at dusk (**Figure 2D**). Control experiments monitoring GFP expression under the control of *Clk-GAL4* confirmed that the dawn-biased proteome does not result from time-dependent fluctuations in APEX2 levels (**Figure 2B**). Principal component analysis (PCA) revealed that experimental condition (Experimental vs. control animals) was the major sources of proteomic variation (PC1, 59.8% of variance), and dawn and dusk samples formed distinct clusters in the experimental group (**Figure 2E**). We noticed that the dawn controls were more variable than the dusk’s. However, experimental replicates showed very high correlation, and Gene Ontology (GO) of cellular component classification confirmed a strong enrichment for plasma membrane and cell-periphery terms, validating our labeling design (**Figures 2F–H**).

### Dawn and dusk proteomes exhibit distinct functional signatures

Of the 956 total proteins identified, approximately half (456) were shared between the two time points of sample collection, while the remainder were specific to either dawn or dusk (**Figure 2D**). Notably, the dawn proteome contained ∼ six times more unique proteins (429) than the dusk proteome (71). This bias might reflect the extensive structural reconstruction the s-LNvs undergo daily^36^. Higher levels of activity-dependent translation could also be a contributor, since most circadian neurons are active at dawn.

Functional annotation revealed distinct biological signatures for each time point (**Figure 2H**). The dawn proteome was significantly enriched for GPCR signaling components, whereas the dusk proteome featured proteins associated with Rho GTPase signaling. This is consistent with previous findings that Rho1 activity peaks at dusk to repress the extension of s-LNv axonal terminals^37^. Interestingly, a recent proteomic analysis of the mammalian suprachiasmatic nucleus (SCN) revealed that Rho GTPase phosphorylation is enriched during the daytime, coinciding with the rest phase of nocturnal mice^38^. Both time points showed enrichment for neuronal regulators (**Figure 2H**). Strikingly, our dataset captured several established regulators of circadian behavior, such as the sodium channel Narrow Abdomen (NA), which as mentioned above regulates excitability in DN1p neurons to guide locomotor behavior output^18,39^, as well as KCC^40^ and HK^41^ (**Figure 2I**). The presence of membrane proteins that are established regulators of circadian behavior underscores the reliability of our platform for discovering novel circadian regulators.

To further refine our search for candidates regulating membrane physiology, we filtered the dataset for "plasma membrane" GO terms, yielding 288 and 156 proteins at dawn and dusk, respectively. Notably, while Rho GTPase signaling was a top-tier term in our Reactome^42,43^ analysis of the total enriched proteome (**Figure 2H**), it was less prevalent when strictly filtering for "plasma membrane" annotations (**Figure 3A**). This is likely due to the classification of many Rho GTPases as cytoplasmic or cytoskeletal components, despite their critical membrane-related function^44^. We compared our proteomic results to a pseudo-bulk analysis of existing clock neuron single-cell RNA-seq (scRNA-seq) data^25^. We compared the transcripts annotated with “plasma membrane” among the top 1,000 transcripts to our plasma membrane proteome and found a significant discordance. Only ∼35% of dawn proteins and ∼18% of dusk proteins had corresponding rhythmic mRNA expression (**Figure S1**). This misalignment mirrors observations in other systems^27,28,45–47^ and might results from the impact of translational lag, selective membrane trafficking, protein degradation or other posttranslational regulations. These results demonstrate that the membrane proteome provides a unique layer of information that cannot be predicted solely by transcriptomics, allowing for the identification of high-priority candidates with membrane expression.

**Figure 3.**
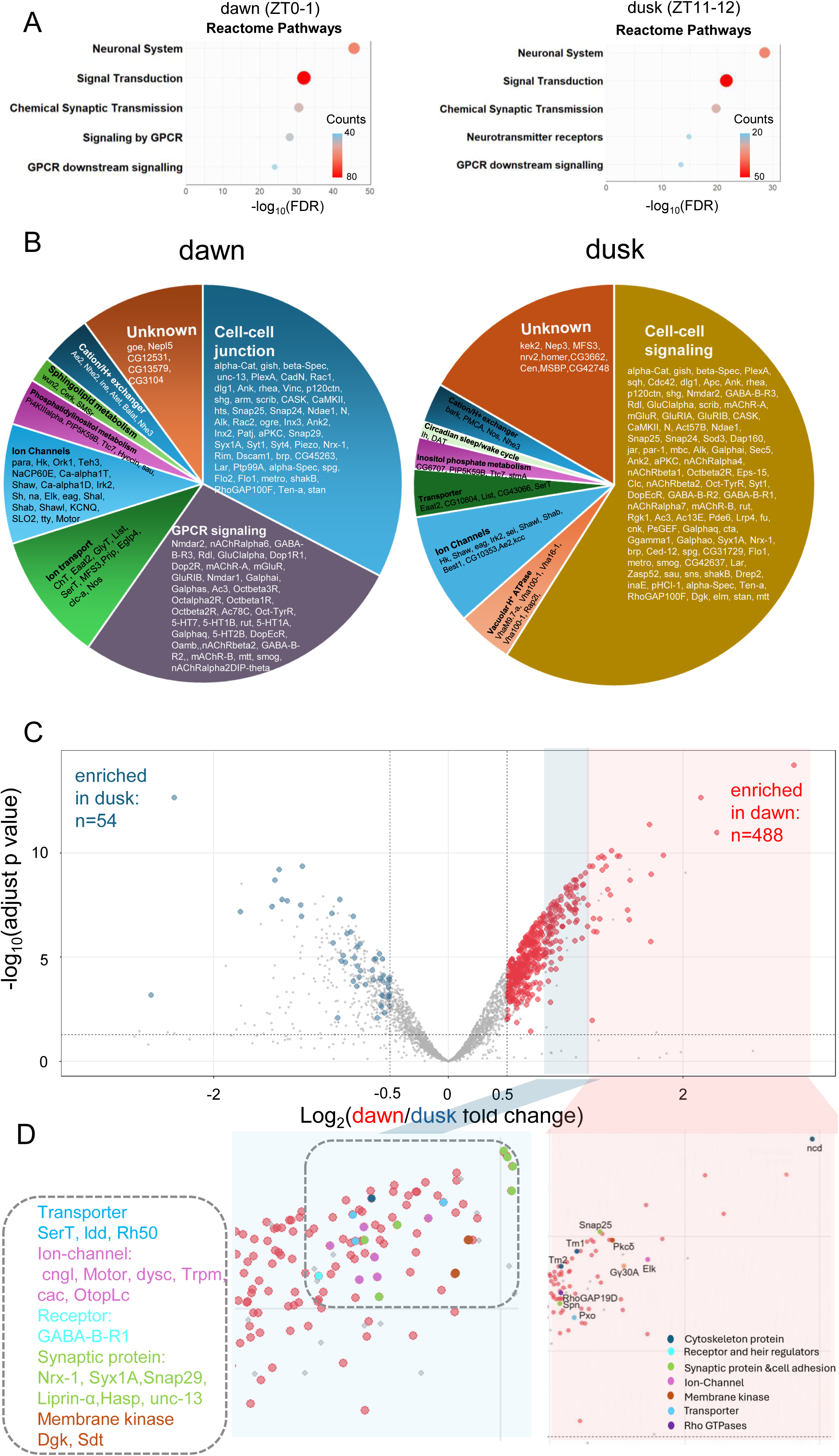
Characterization of the plasma membrane-enriched proteome in clock neurons. (A) Top 5 Reactome pathways for the clock neuron membrane proteome at two time points. (B) Comparison of plasma membrane protein clusters enriched in clock neurons at dawn (left, ZT0-1) and dusk (right, ZT11-12). Markov clustering of enriched plasma membrane proteins at each time point was performed using protein-protein interaction scores from the STRING database. Note that only representative proteins in each cluster are shown for clarity; the full dataset is available in Data S2. (C) Volcano plot displaying the differential abundance of cell-membrane proteins (CMPs) at dawn (ZT1) compared with dusk (ZT11). Each red/blue point represents an individual protein that are enriched at either dawn(red) or dusk(blue) after validated by their negative controls, while each grey diamond indicates non-significant protein. Proteins significantly downregulated at dawn compared with dusk (log_2_[fold change] < −0.5 and adjusted p-value < 0.05) are shown in blue, while proteins significantly upregulated at dawn compared with dusk (log2[fold change] > 0.5 and adjusted p-value < 0.05) are shown in red. (D) Zoom-in of dawn-enriched proteins with log2FC>1(red background) and 0.7<log2FC<1 (blue background) proteins are color-coded based on their molecular functions. See also **Figure S1**, **Table S1**, and **Data S2**.

To further explore whether the enriched membrane proteins form distinctive functional blocks, we employed a Markov Cluster Algorithm to visualize the protein-protein interaction (PPI) network of the enriched proteins (**Figure 3B; Data S2**). This unsupervised clustering identified a distinct proteome landscape when comparing dawn and dusk, with “cell-cell junction”, "GPCR signaling", and “lipid metabolism” modules at dawn and a "circadian/sleep-wake" module at dusk, suggesting that different molecular mechanisms govern morning and evening neural activity and function in the clock neurons.

In addition to exploring the dawn and dusk proteomes separately, TMT labeling enabled a quantitative comparison of protein levels between the two time points (**Figures 3C–D**). We found that 488 proteins were more abundant at dawn than at dusk, whereas 54 proteins exhibited the opposite pattern. Interestingly, the majority of receptors and ion channels were enriched at dawn (**Figure 3D**)—a global upregulation of these cell-membrane proteins that is consistent with the fact that most clock neurons (except for the LNds) exhibit high level of Ca^2+^ signals at dawn^20^. Among the proteins with higher abondance at dawn, we identified membrane proteins either known to be expressed in clock neurons (e.g., nAchRβ2^48^,MTT^49^) or implicated in regulating circadian rhythms (e.g., NA^18,39^, Shaw^19,50,51^) (summarized in **Table S1**).

### A proteome-informed genetic screen identifies novel regulators of circadian rhythms

A significant proportion of the membrane proteins identified in our clock proteome have not been previously characterized in the context of circadian biology (**Figures 3C-D; Tables S2-3**). To determine whether these proteins regulate clock function, we conducted a targeted genetic screen using RNA interference (RNAi; **Figure 4A; Table S2**) and null mutants (**Figure S3A; Table S3**). We tested flies under standard 12h/12h light/dark (LD) cycle followed by constant darkness (DD). Of the 110 candidates tested, 29 (26%) exhibited significantly reduced rhythmicity or an extended period length (**Figures 4A, S3; Tables S2-3**). For example, silencing the synaptic exocytosis-related proteins UNC-13^52^ and SYX1A^53^ specifically in clock-expressing tissues via *tim-GAL4* or only in clock neurons with *Clk-Gal4* severely reduced rhythmicity under DD (**Figures 4B-C, and S2A-B**), Similarly, proteins associated with synaptic vesicle cycling exhibit rhythmic expression in the mammalian SCN, and pharmacologically blocking synaptic vesicle cycling dampens molecular clock oscillations under DD^54^.. Interestingly, under LD, SYX1A downregulation had little impact on circadian behavior under LD, but UNC-13 reduction had a more severe effect under these conditions, particularly when driven by *Clk-GAL4*. We also found that candidates encoding Gap junctions (Ogre (Inx1) and Inx2) are required for robust rhythmicity under DD (**Table S2**). These two proteins had previously been found to impact circadian behavior period in DD ^55^. Similarly, GABA receptor (GABA-B-R2) null mutants severely disrupted circadian behavior under DD, with a high level of arrhythmicity (**Figures S3 B-C**). In contrast, knockdown of *Motor* (*CG3638*) - which encodes a Ca^2+-^activated chloride channel - using two independent RNAi constructs driven by either *tim-GAL4* or *Clk-GAL4* led to a significant lengthening of the circadian period (**Figures 4D-E, and S2C**). These results demonstrate that our membrane proteome is a rich resource for identifying novel molecular mechanisms that govern circadian behavior.

**Figure 4.**
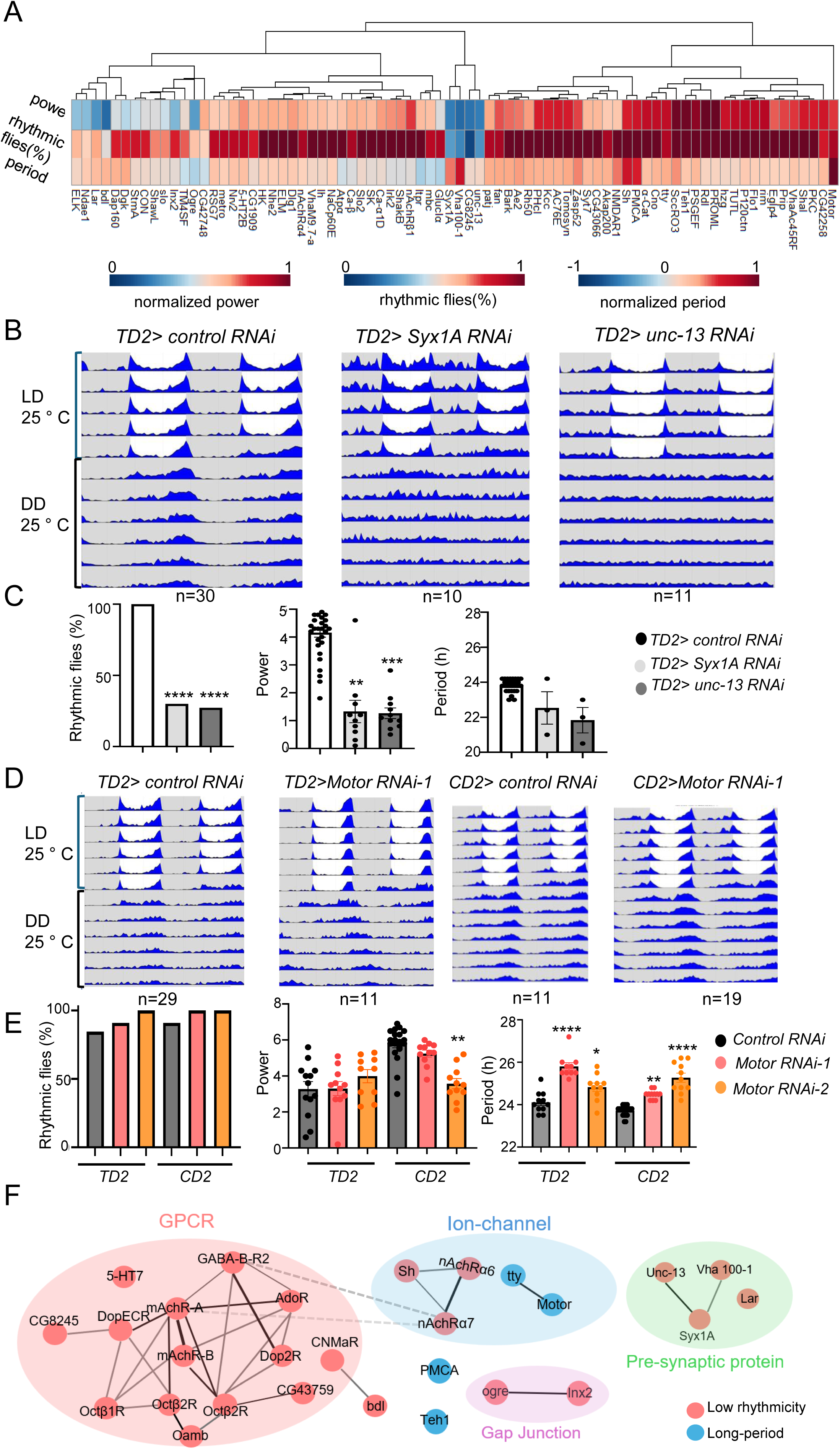
A proteome-guided genetic screen identifies novel regulators of circadian rhythmicity. (A) Heatmap summary of rhythmicity percentage, power, and period length in DD following RNAi-mediated knockdown or null mutation of candidate genes. Data are normalized as follows: rhythmic power: x_i_/x_max_, and period length is normalized as: (X_i_-X_control_)/( X_max_-X_control_), while the percentage of rhythmic flies is presented as original (raw) values alongside respective controls for direct comparison. Each candidate was tested with n >=10 flies. (B) Knockdown of *Syx1A* and *unc-13* via *tim-GAL4* (TD2) results in severely reduced rhythms. Double plotted actograms (top) and daily average locomotor activity in DD are shown for control (mCherry RNAi), *Syx1A* RNAi, and *unc-13* RNAi. (C) Quantification of behavior for the genotypes in (B). For period length, only rhythmic flies were included. A Fisher’s exact test was used to determine the statistical significance of differences in percentage of rhythmic flies, while a Kruskal–Wallis test followed by Dunn’s multiple-comparisons test was used to determine the significance of changes in rhythmic power and period length. ****p < 0.0001. *** p < 0.001, ** p < 0.01. (D) Knockdown of *Motor* (*CG3638*) leads to a significantly lengthened period in DD. (E) Same as in C, but quantification of rhythmic parameters for *Motor* knockdown. (F) Markov clustering of screen hits was performed with protein-protein interaction scores from the STRING (search tool for the retrieval of interacting genes/proteins) database^90^. Gray/black lines denote protein-protein interactions, dash lines indicate weak interaction. See also **Figures S2-3, Tables S2-3.**

Protein-protein interaction (PPI) network analysis revealed that the genes identified in our genetic screen form distinct functional clusters (**Figure 4F**). Specifically, most GPCRs and gap junctions were required to maintain behavioral rhythmicity under DD, whereas chloride channels (Motor and TTY) were essential for maintaining proper period length. These findings suggest that cell-membrane proteins (CMPs) organize into specialized networks to regulate distinct behavioral tasks.

### GPCR Octβ1R is essential for behavioral adaptation to long photoperiods

We next investigated whether some of our candidates are involved in seasonal behavior adaptation to different photoperiods. In wild-type *Drosophila*, the circadian network adjusts the phase of morning and evening activity to the length of the day^56^. For example, under summer-like conditions, the evening peak of locomotor activity shifts away from the morning peak. We first used a summer-like 18:6 LD cycle, under which evening activity typically precedes the lights-off transition by approximately 2–3 hours, whereas it closely coincides with lights-off under standard 12:12 L:D cycles **(Figures 5A–B)**. We performed a genetic screen of our candidates under this long photoperiod cycle and identified one candidate with an advanced evening phase, Octβ1R, which encodes an octopamine G protein-coupled receptor (GPCR, **Figures 5C-D**). Evening activity in *Oct*β*1R^ko^* mutant flies^57^ peaked 4–5 hours prior to lights-off, which is ∼2 hours earlier than in controls. In other words, these mutant flies did not adapt properly to the long photoperiod. *Oct*β*1R^ko^* flies also exhibited reduced behavioral rhythmicity under DD (**Figure S3B-C**), but their morning and evening activity peak of activity were normally phased under standard 12:12 L:D conditions (**Figure S5C**).

**Figure 5.**
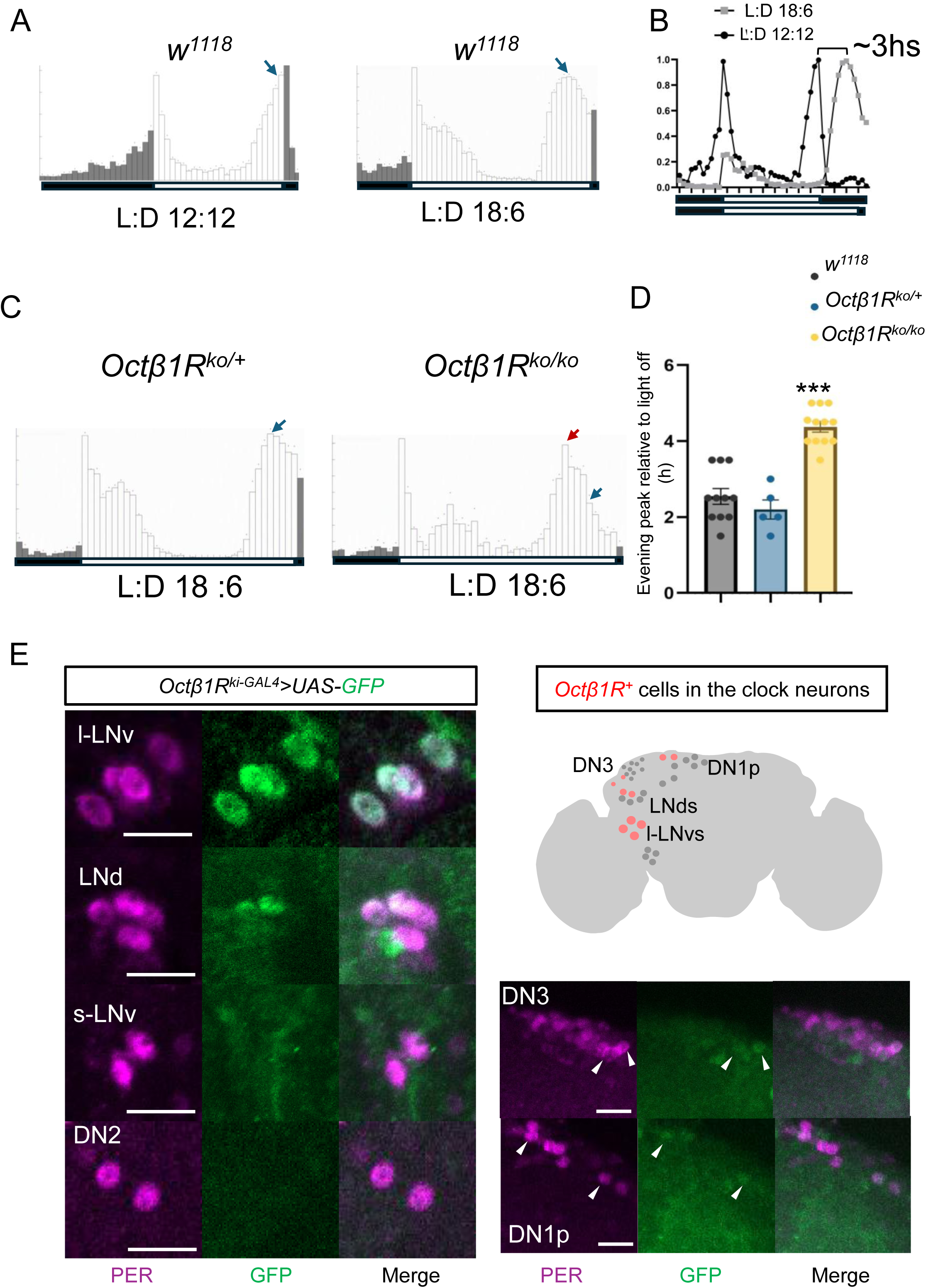
Octβ1R is required for behavioral adaptation to long photoperiod. (A) Daily average locomotor activity of *w^1118^* flies under normal photoperiod (left) and long photoperiod (summer-like,right). (B): Normalized activity shows a ∼3-hour delay in the evening peak under long photoperiod compared with the normal photoperiod (L:D 12:12). (C) Representative locomotor activity of *Oct*β*1R^ko/+^*(n=11) and *Oct*β*1R^ko^* (n=11) under long photoperiod. *Oct*β*1R* mutants exhibit an evening peak advance compared with *w^11^*^18^. (D) Quantification of the evening peak phase, expressed as the number of hours preceding the light-off transition. Each dot represents an independent experiment (n=8–10 flies per experiment). ****p < 0.001, one-way ANOVA with Tukey’spost-hoc test. (E) Expression of a cytoplasmic GFP driven by *Oct*β*1R^kiI-GAL^*^4^ co-labeled with anti-PER. As shown in the schematic, *Oct*β*1R* us highly expressed in l-LNvs and weakly expressed in DN1ps, DN3 and LNds, but not in s-LNvs. Scale bars, 20 μm. See also **Figures S3 and S5**.

Previous single-cell RNA-seq data from *Drosophila* clock neurons indicated that *Oct*β*1R* is expressed in a subset of pacemaker cells—including the DN1ps, DN2s, and E-oscillators (LNds)^24^. We mapped its spatial distribution using an *Oct*β*1R -GAL4* knock-in (ki) line^57^. Confocal analysis confirmed weak expression in the DN1ps and LNds, alongside robust expression in the large ventral lateral neurons (l-LNvs) and weak expression in the DN3 neurons (**Figure 5E**). Given that the l-LNvs are established regulators of evening peak phase under long photoperiods via PDF signaling^58^, these findings indicate that Octβ1R acts within the clock network to coordinate behavioral adaptation to summer-like conditions.

### Piezo is essential for behavioral adaptation to short photoperiods

We also performed a genetic screen with our candidates under short photoperiods (winter-like conditions, 6:18 LD, 25°C). Under such condition, the morning peak of activity precedes the light-on transition by approximately 4.5 hours in wild-type flies (**Figure 6A**).

**Figure 6.**
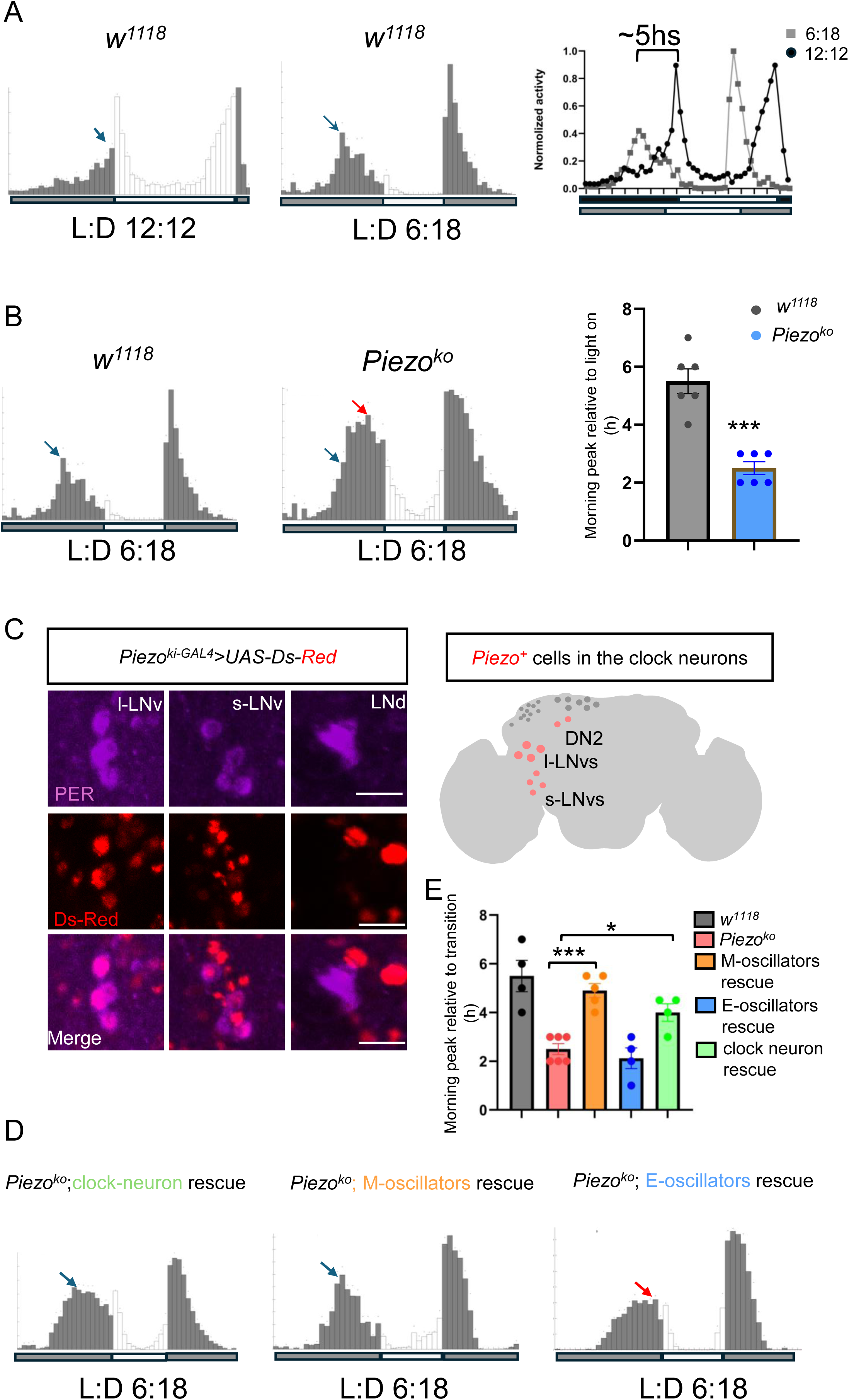
Piezo is required for behavioral adaptation to short photoperiod. (A) Daily average locomotor activity of *w^11^*^18^ flies under short photoperiod (winter-like, middle) and normal photoperiod (left). Right: Normalized activity shows a ∼5-hour advance in the morning peak under short photoperiods. (B) Representative locomotor activity of *w^11^*^18^ (n=11) and *Piezo^ko^* (n=11) under short photoperiod. *Piezo* mutants exhibit a morning peak delay compared with *w^11^*^18^. Right: Quantification of the morning peak phase, expressed as the number of hours preceding the light-on transition. Each dot represents an independent experiment (n=8–10 flies per experiment). ***p < 0.001, Mann–Whitney U test. (C) Expression of a nuclear ds-Red marker driven by *Piezo^ki-GAL4^*co-labeled with anti-PER. As shown in the schematic, Piezo expression is found in the DN2s and LNvs, but not in the LNds. Scale bars, 20 μm (D) Restoration of wild-type *Piezo* in LNvs, but not LNds, rescues the mutant phenotype. (E) Quantification of morning peak phase for the indicated rescue genotypes. Kruskal–Wallis test followed by Dunn’s multiple-comparisons test, ***p < 0.001, *p < 0.05,. See also **Figures S3, S4 and S5**.

We found that disruption of 6 candidates caused a shift in morning phase under 6:18 LD cycles (**Figures 6B, and S4**,). Among these candidates, we identified Piezo^30^, a mechanosensitive ion channel, as a key regulator of photoperiodic adaptation. Compared with *w^11^*^18^ control flies, *Piezo* knockout (*ko*) mutants exhibited a significantly delayed morning activity peak, occurring only 1.5–2 hours before lights-on under short photoperiods (**Figure 6B**). In contrast, *Piezo^ko^* mutants exhibited a normal morning activity peak under standard 12: 12 LD conditions, similar to the wild-type controls. (**Figure S5B**) These mutants thus shift more strongly their morning activity peak in response to short photoperiod than wild-type controls. *Piezo^ko^* flies also exhibit normal rhythmic behavior under DD (**Figures S3B-C**).

Using a *Piezo-GAL4* knock-in line^59^, we determined that *Piezo* is expressed in a small subset of clock neurons, specifically the large and small ventral lateral neurons (l-LNvs and s-LNvs) and the DN2s (**Figure 6C**). Consistent with this expression pattern, restoring wild-type *Piezo* expression specifically in the LNvs (M-oscillators) rescued the morning peak defects in *Piezo^ko^* flies, whereas restoration in the LNds (E-oscillators, by *GMR 16C05-GAL4*^60^) failed to rescue the phenotype (**Figures 6D–E**). Restoration of Piezo in l-LNvs did not rescue the morning phase phenotype (**Figure S5G**), indicating that Piezo function in s-LNvs to modulate morning activity peak under short photoperiod.

### Piezo mediates clock neuron activity and the daily structural plasticity of s-LNv axonal terminals

To determine whether Piezo directly regulates clock neuron activity, we utilized CaMPARI2^61^, a photoconvertible Ca^2+^ indicator, to monitor real-time Ca^2+^ dynamics across circadian pacemaker subpopulations **(Figure 7A)**. Consistent with previous GCaMP-based studies^17,20^, wild-type M-oscillators (s-LNvs) exhibited anti-phase activity relative to E-oscillators (LNds) under a standard 12:12 LD cycle. This temporal relationship was completely abolished in *per*^01^ null mutants, validating our approach **(Figures 7B–C)**.

**Figure 7.**
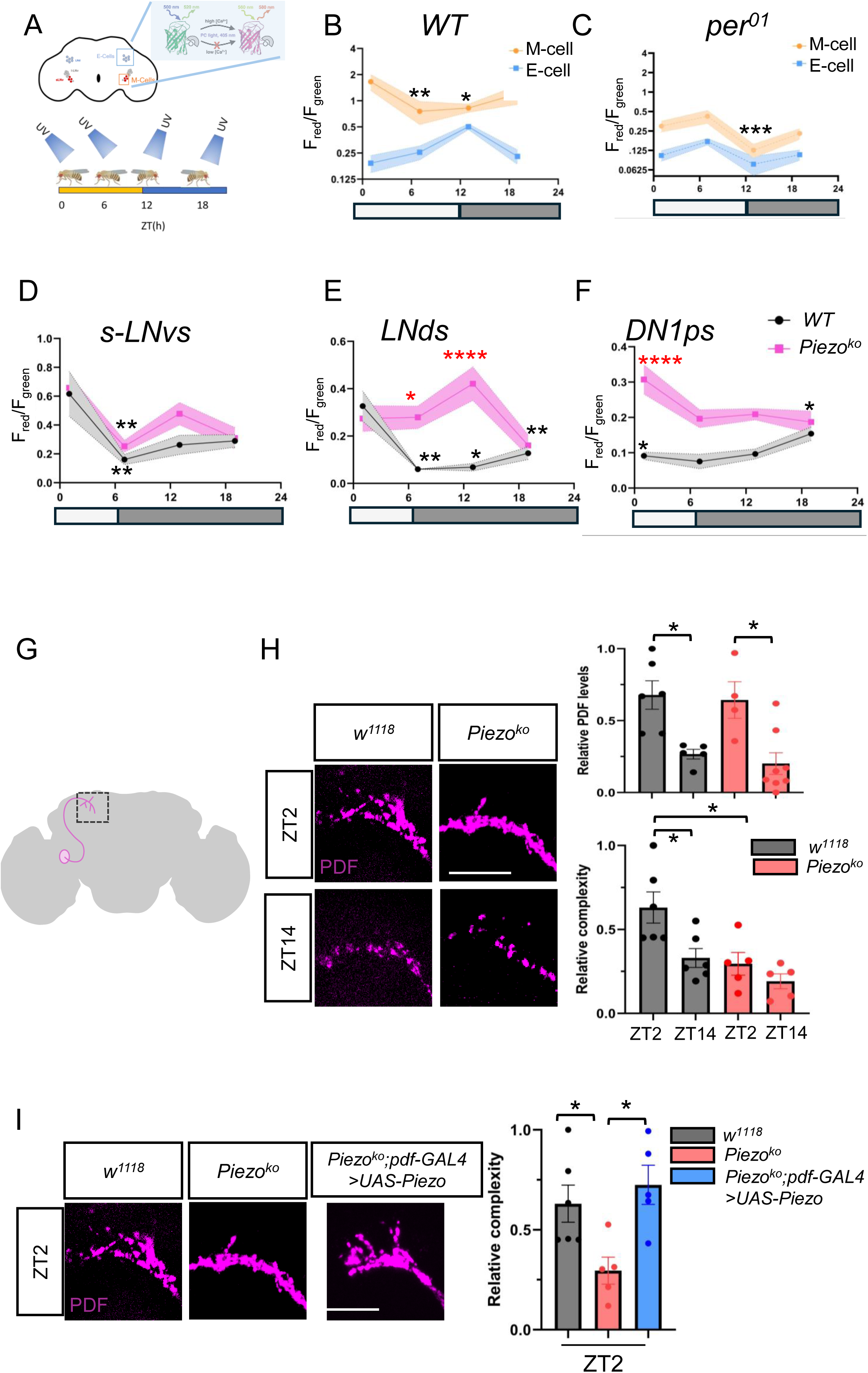
Piezo regulates activity phase of clock neurons and the circadian structural plasticity of s-LNv dorsal projections. (A) Schematic representation of *in vivo* Ca^2+^ imaging experiment using CaMPARI2. Male flies expressing CaMPARI2 in clock neurons were sampled at indicated time points. (B-C) Quantification of the normalized F_red_/F_green_ ratio of CaMPARI2 in the M (morning, sLNvs) and E(evening, LNds) clock neurons in the WT (B) and *per*^01^ (C) under 12:12 LD, error bars, SEM. ***P < 0.001, **P < 0.01 and * P < 0.05, two-way ANOVA with Tukey’s post hoc test. Asterisks represent significant differences relative to the peak value within the same genotype. (D-F) Quantification of the normalized F_red_/F_green_ ratio of CaMPARI2 in different subsets of clock neurons in the WT and *Piezo^ko^* under short photoperiod, error bars, SEM. ****P < 0.001, ***P < 0.001, **P < 0.01 and * P < 0.05, two-way ANOVA with Tukey’s post hoc test. Red asterisks indicate significant differences between WT and mutant flies, whereas black asterisks represent significant differences relative to the peak value within the same genotype. (G) Schematic representation of s-LNv branched terminals in the dorsal brain. (H) Left panels. Representative confocal images of s-LNv projections from control and *Piezo^ko^* mutants dissected at ZT2 and ZT14. Endogenous PDF staining (magenta) labels the axonal projections to visualize and quantify s-LNv axonal terminal morphology. Scale bars, 20 μm. Right panels. Quantification reveals a loss of plasticity rhythms in the mutant flies, with a significant reduction in the structural complexity of s-LNv arborizations relative to controls at ZT2. PDF abundance rhythm is not impaired by loss of Piezo. Each dot represents the terminals of a single hemibrain. (I) The retracted axonal phenotype in Piezo mutants at ZT2 is rescued by expression of wild-type PIEZO in the LNvs via *pdf-GAL4*. *p < 0.05, Kruskal–Wallis test followed by Dunn’s multiple-comparisons test. See also **Figures S5**.

When evaluated under short-photoperiod conditions, wild-type M- and E-oscillators both peaked around ZT1, whereas DN1p neurons peaked at ZT18. In *Piezo* mutant flies, M-oscillator peak timing remained unchanged at ZT1; however, we observed a striking 12-hour phase shift in E-oscillators and a 6-hour phase delay in DN1p neurons (**Figures 7D–F**). Importantly, the DN1ps include morning-activity promoting neurons controlled by the s-LNvs in wild-type flies^62^. The delayed DN1p activity thus likely explains the delayed morning-peak under short photoperiod. Given that Piezo is not expressed in DN1ps or in E-oscillators, the altered phase in neural activity rhythms in these neurons implies that Piezo acts non-cell-autonomously to coordinate network-level timing.

Under standard 12:12 LD conditions, *Piezo^ko^* flies displayed significantly dampened rhythms in Ca^2+^ signals in DN1ps and LNds compared with controls, but the phase of these rhythms was unchanged, thus explaining the proper timing of the M and E peaks of activity (**Figures S5 E-F**). However, the phase of s-LNv oscillation was unexpectedly altered under 12:12 LD cycle while it was not altered under short photoperiod (**Figure S5D**). This indicates that photoperiod length determines the impact of Piezo on the timing of rhythmic activity in the circadian neuronal network. Taken together with our behavioral rescue data, these physiological findings demonstrate that Piezo is required for seasonal behavioral adaptation by fine-tuning neural activity dynamics across interconnected clock circuits, including the LNds and DN1ps.

The s-LNvs undergo daily morphological remodeling that include changes in synaptic connectivity^63^. This restructuring is critical for seasonal timing^64^. Intriguingly, Piezo was recently found to regulate dendrite targeting in the developing projection neurons in the olfactory system^65^. We therefore decided to examine whether Piezo is required for M-oscillators’ structural plasticity. We visualized s-LNv morphology using anti-PDF staining (**Figures 7G-H**). In control flies under LD conditions, the s-LNv axonal termini exhibited a clear circadian rhythm in structural complexity as previously reported^36,37^: they were maximally spread at dawn (ZT2) and significantly retracted at dusk (ZT14), a transition confirmed by 3D spread quantification (**Figure 7H**). In contrast, the axonal terminals of *Piezo* mutants showed a significant reduction in branching complexity at ZT2, appearing "locked" in a retracted, dusk-like state at both time points (**Figure 7H**). Importantly, the rhythmic expression and distribution of the PDF neuropeptide itself were not affected, suggesting that the defect is structural rather than a failure of PDF production, transport or release. Furthermore, restoration of *Piezo* expression in the M-oscillators using *pdf-GAL4* was sufficient to rescue these structural deficits (**Figure 7I**). Taken together, these data indicate that Piezo has a critical role in regulating the daily structural plasticity of s-LNvs required for proper circadian adaptation to different photoperiods.

## Discussion

Despite the fundamental role of membrane proteins in cell-cell communication and neuronal development, systematically identifying these proteins within specific neuronal cell types remains a formidable challenge. Traditional biochemical fractionation often suffers from significant contamination by nuclear and mitochondrial membranes and, crucially, lacks the spatial resolution necessary to distinguish specific intermingled neuronal populations. This challenge is particularly daunting for the circadian neural network of *Drosophila*, which is composed of only 240 brain neurons, out of about 150,000, express core clock genes. They are dispersed in a very small brain. Here, by combining cell-type-specific proximity labeling (APEX2) with quantitative TMT-based mass spectrometry, we have established a robust pipeline for labeling the membrane-associated proteome of *Drosophila* clock neurons *in situ,* isolating these proteins and identifying them. Our Gene Ontology (GO) analysis confirmed a high degree of enrichment for plasma membrane proteins, validating the spatial specificity of our approach. Moreover, we identified known membrane regulators of circadian rhythms, such as *NA*, *KCC*, *HK* and *SHAL*^18,19,41,66^. However, we noticed the surprising absence of PDF-receptor (*PDFR*), which is expressed in several groups of circadian neurons^67^. This could reflect a particularly low level of *PDFR* expression in these neurons, or perhaps this protein was not accessible to the APEX2 enzyme.

A striking finding of our study is the strong bias for protein enrichment at dawn (ZT1) vs dusk (ZT11). This temporal asymmetry may be driven by two non-exclusive mechanisms. First, given that the majority of the clock network (with the exception of E-oscillators) is preferentially electrically active at dawn^17^, this enrichment may reflect activity-dependent protein translation and trafficking. Second, this bias might be related to the well-documented structural plasticity of pacemaker neurons^36,37,68,69^. Specifically, M-oscillators undergo a daily expansion of their axonal terminals at dawn; the increased protein diversity we observed may be a direct consequence of this expanded surface area or, more intriguingly, may represent the molecular machinery required to drive this structural remodeling. Indeed, loss of *Piezo* compromises this restructuring. Future studies utilizing specific antibodies against these dawn-enriched candidates will be essential to distinguish between these possibilities. Intriguingly, while abundance remains constant, phosphorylation of several membrane proteins in the mammalian SCN also exhibits a dawn peak^38^. This indicates that coupling neural activity with membrane protein dynamics may be an evolutionarily conserved feature of circadian regulation in the central pacemaker. It would thus be important to also explore the *Drosophila* phospho-proteome of circadian plasma membrane proteins in the future.

The reliability of our proteomic dataset is further supported, as discussed above, by the robust detection of established circadian regulators. This provided us with a high-confidence foundation for our proteome-informed genetic screen, which successfully identified numerous novel regulators of circadian behavior, impacting rhythmicity, period length, and environmental adaptation. By bridging spatiotemporal proteomic "snapshot" and functional genetics, we have demonstrated that the membrane proteome is a critical, yet previously under-explored, dynamic regulator of circadian behavior. In addition, this dataset will serve as a valuable resource for the community to investigate other facets of clock neuron physiology.

Seasonal adaptation—the ability to adjust physiology and behavior to changing photoperiods—is a vital survival strategy across species. While this phenomenon can be observed in *Drosophila,* for example at the behavioral level^56,58,64,70,71^, the molecular mechanisms that enable the clock network to "perceive" and adapt to these changes have remained unclear. We identified the mechanosensitive ion channel Piezo as a critical contributor to behavioral adaptation to short photoperiods. Interestingly, *Piezo* mutants exhibit an exaggerated, "over-adapted" phase change under short photoperiods. This indicates that Piezo normally act as a regulatory "brake" under winter-like conditions. Surprisingly, under short photoperiod, loss of *Piezo* does not alter the phase of the M-oscillators themselves but disrupts the timing of downstream circadian nodes. Consistent with this, Piezo appears to govern the magnitude of structural remodeling in s-LNv axonal terminals, a well-established driver of seasonal behavioral plasticity. Given Piezo’s canonical role in mechanotransduction^30^, it might sense the physical expansion of the s-LNv axonal arbor, providing crucial feedback to the molecular clock to coordinate network-wide timing and behavioral output in response to environmental cues.

While our results indicate that Piezo functions in s-LNvs to regulate neural activity in the circadian neural network and adaptation to short photoperiod through rhythmic structural reorganization of their projections, Octβ1R likely controls adaptation to long photoperiod through a distinct mechanism, as it is predominantly expressed in Evening-controlling neurons: the l-LNvs and, more weakly, the LNds. It is however not expressed in the s-LNvs, suggesting that adaptation to long or short photoperiods relies on specific network mechanisms and distinct neurons.

In summary, our work identifies numerous novel membrane-associated proteins enriched in *Drosophila* circadian neurons, and our screens validate the critical importance of this unique dataset. Combined with the remarkable power of *Drosophila* genetics and neural imaging, we anticipate that this resource will lead to a comprehensive molecular understanding of the circadian neural network and the mechanisms by which it controls various circadian behaviors and their specific temporal patterns.

## Materials and Methods

### Fly strains and husbandry

Flies were reared in plastic vials with cotton plugs containing standard yeast-enriched fly food. Vials were kept in an environmentally controlled room, where they were entrained to a 12hr:12hr light-dark (LD) cycle at constant 25°C. The following fly lines were used in this study: *pdf-Gal4* (on 2^nd^ chromosome)^8^, *pdf-Gal4* (on 3^rd^ chromosome)^72^, *clk-856-Gal4*^34^, *piezo^ki^-Gal4*^59^, *tim-Gal4, UAS-dicer2 (TD2)*^73^, *GMR-16C05-Gal4*^60^, *GMR-78G04-Gal4*^60^*, UAS-CD8:APEX2*^32^, *UAS-dsRed*^74^, *Oct*β*1R^ko^* ^57^, *Oct*β*1R^KI^-GAL4*^57^*, Piezo^ko^* ^30^, *UAS-Piezo-EGFP*^30^, *UAS-mCherry RNAi* (BDSC#35785), UAS-EGFP (BDSC#5428), UAS-CaMPARI2 (BDSC#78318), *UAS-syx1A RNAi* (BDSC#25811), *UAS-unc-13 RNAi* (BDSC#29548), *UAS-Motor RNAi-1* (VDRC#102444), UAS-Motor RNAi-2 (VDRC#6914), The RNAi flies used for genetic screen are listed in **Tables S2-3**.

### Cell membrane protein biotinylation in fly brains

Cell-membrane biotinylation was performed using a modified proximity-labeling protocol^26,65^. Briefly, 3- to 5-day-old adult flies expressing UAS-CD8:APEX2 under the control of *pdf-GAL4* or *Clk-GAL4* were entrained under 12:12 light:dark cycles at 25°C (LD 25) for 5 days. For each biological replicate, approximately 800 fly brains were dissected in ice-cold extracellular saline^75^ and collected in 0.5 mL low-protein-binding tubes (Eppendorf). Following dissection, brains were washed thoroughly with fresh saline to remove fat bodies and debris. Samples were pre-incubated with 100 µM biotin-phenol (BP; Sigma-Aldrich) in saline for 1 hour at 4°C with gentle rotation. To initiate proximity labeling, brains were incubated with 1 mM H_2_O_2_ (Thermo Fisher Scientific) for exactly 5 minutes at 4°C. The reaction was immediately quenched by five rapid washes with quenching buffer (10 mM sodium ascorbate, 5 mM Trolox, and 10 mM sodium azide in PBS). For histological validation, brains were fixed in 4% paraformaldehyde (PFA) immediately after quenching. For proteomic analysis, biotinylated brains were snap-frozen in liquid nitrogen and stored at –80°C until further processing. In total, approximately 6,400 brains were processed to obtain the required input for TMT proteomic experiment.

### Lysis of fly brains and protein extraction

Frozen fly brains were homogenized in 40 µL of high-SDS RIPA buffer (50 mM Tris-HCl, 150 mM NaCl, 1% SDS, 0.5% sodium deoxycholate, 1% Triton X-100, 1× protease inhibitor cocktail [Sigma-Aldrich, P8849], and 1 mM phenylmethylsulfonyl fluoride [PMSF]). Homogenization was performed on ice using disposable motorized pestles. For each experimental group, lysates were pooled into a single tube, and the volume was adjusted to 300 µL with high-SDS RIPA buffer. Samples were vortexed briefly and subjected to two rounds of sonication (10 seconds each; Virtis 100). Next, 1.2 mL of SDS-free RIPA buffer (50 mM Tris-HCl [pH 8.0], 150 mM NaCl, 0.5% sodium deoxycholate, 1% Triton X-100, 1× protease inhibitor cocktail, and 1 mM PMSF) was added to each sample. The lysates were rotated for 1 hour at 4°C to ensure complete solubilization. Subsequently, samples were transferred to 3.5 mL ultracentrifuge tubes (Beckman Coulter) containing 200 µL of standard RIPA buffer (Thermo Fisher Scientific, 88900) and centrifuged at 100,000 × g for 30 minutes at 4°C. The supernatant (approx. 1.5 mL) was carefully collected and kept on ice for subsequent affinity enrichment.

### Affinity enrichment of biotinylated proteins

To enrich biotinylated proteins, 400 µL of streptavidin-coated magnetic beads (Pierce, 88817) were washed twice with 1 mL of RIPA buffer. The pre-washed beads were added to each of the post-ultracentrifugation brain lysates and incubated overnight at 4°C with gentle rotation. The following day, the beads were washed sequentially to minimize non-specific binding: twice with 1 mL of RIPA buffer, once with 1 mL of 1 M KCl, once with 1 mL of 0.1 M Na_2_CO_3_, once with 1 mL of 2 M urea in 10 mM Tris-HCl (pH 8.0), and finally twice with 1 mL of RIPA buffer. To elute the captured proteins, the beads were resuspended in 2% SDS and boiled at 95°C for 10 minutes. The eluate was then diluted fourfold with Milli-Q water and vortexed for 10 minutes. The supernatant containing the enriched membrane proteome was collected and processed for downstream TMT labeling.

### Western blots

To analyze the enriched protein fraction, biotinylated proteins were eluted from streptavidin beads by adding 20 µL of elution buffer (2× Laemmli sample buffer, 20 mM DTT, and 2 mM biotin) and boiled at 95°C for 10 minutes. Proteins were resolved on 4–20% Mini-PROTEAN TGX Stain-Free Protein Gels (Bio-Rad Laboratories) and transferred to nitrocellulose membranes (Thermo Fisher Scientific) using a Trans-Blot® SD Semi-Dry Transfer Cell (Bio-Rad). All wash and incubation steps were performed on an orbital shaker at room temperature. Membranes were blocked with Intercept Blocking Buffer (LI-COR) for 1 hour, followed by incubation with primary antibodies for 1 hour at room temperature or overnight at 4°C. Membranes were washed three times for 5 minutes each in TBST (Tris-buffered saline with 0.2% Tween 20; Thermo Fisher Scientific). Subsequently, membranes were incubated with fluorescent-conjugated secondary antibodies (LI-COR) for 1 hour, followed by four additional 5-minute washes with TBST. Protein bands were visualized using an Odyssey DLx Imaging System (LI-COR).

Primary antibodies used in this study were: mouse anti-tubulin (1:2,000; ab8224, Abcam) and mouse anti-Spectrin (1:5,000; 3A9, Developmental Studies Hybridoma Bank). The secondary antibody was IRDye 800CW (LI-COR, cat # 925-32211) used at 1:10,000. To detect total biotinylated proteins, membranes were incubated with fluorescent-conjugated streptavidin (Thermo Fisher Scientific) instead of antibodies.

### TMT labeling and fractionation of peptides

Biotinylated protein solution samples were flash-frozen on dry ice and dried in a SpeedVac. Then, 23 µL of 2× lysis buffer (10% SDS, 100 mM TEAB) was added, and samples were processed using the S-Trap™ Micro Spin digestion protocol (PROTIFI). Proteins were reduced with 4 mM tris(2-carboxyethyl) phosphine (TCEP, Thermo Scientific) and alkylated with 20 mM iodoacetamide (Millipore Sigma). After overnight digestion with trypsin at 37 °C, peptides were eluted sequentially with 40 µL of 50 mM triethylammonium bicarbonate (TEAB), 40 µL of 0.2% formic acid (FA), and 40 µL of 50% acetonitrile (ACN), with a 1-min spin at 4,000 rpm after each elution. The three eluates were pooled, dried in a SpeedVac, and reconstituted in 100 µL of 100 mM TEAB. Peptide concentration was measured, and equal amounts from each condition were transferred to a new tube for labeling with the TMT10plex Mass Tag Labeling Kit (Thermo Scientific, lot number YL380292) according to the manufacturer’s instructions. A pooled control sample was labeled with the TMT10-130C tag. Each reaction was quenched with 8 µL of 5% hydroxylamine (Thermo Scientific) for 15 min, after which equal amounts of each sample were combined and dried in a SpeedVac. The combined labeled mixture was dissolved in 300 µL of 0.1% trifluoroacetic acid (TFA) and fractionated using the Pierce High pH Reversed-Phase Peptide Fractionation Kit (Thermo Scientific) according to the manufacturer’s protocol. Each fraction was evaporated to dryness in a SpeedVac and reconstituted in 20 µL of 0.1% FA in 5% ACN. Samples were then centrifuged at 16,000 rpm for 16 min, and 18 µL of the supernatant was transferred to non-binding HPLC vials.

### Mass spectrometry data processing

Data was acquired using a Thermo Scientific Vanquish Neo UHPLC coupled to an Orbitrap Ascend Tribrid (Thermo Fisher Scientific, Waltham, MA) mass spectrometer. Peptides were separated by direct injection using an Aurora Ion Optics C18 60cm x 75μm, 1.7 μm 100 Å analytical column, using an aqueous mobile phase (A) of water and 0.1% FA and an organic mobile phase (B) of 80% ACN and 0.1% FA. A 2 µL sample was injected at a maximum 1300 bar pressure on an analytical column kept at 45°C, followed by a gradient elution with 7-28% in 10min, 28-50% in 20min, 55-99% in 10min wash at a flow rate of 250 nL/min over 135 min separation (total runtime: 145 min). Mass spectra were acquired over m/z 400-1600 Da with a resolution of 120,000. Tandem mass spectra were acquired using data-dependent acquisition with an isolation window of 1.2 Da, HCD collision energy of 36%, orbitrap resolution of 30,000 (m/z 200), maximum injection time of 100ms, and a 200% AGC.Liquid chromatography and mass spectrometry

### Proteomic data analysis

Raw files were searched using the Comet search engine (ver. 2019.01.5)^76^with the UniProt fruit fly (*Drosophila melanogaster*) proteome database (downloaded 10/01/2024) with contaminants and reverse decoy sequences appended. Precursor error tolerance was 50 p.p.m. and fragment error tolerance was 0.02 Da. Static modifications included Cys carboxyamidomethylation (+57.0215) and TMT (+229.1629) on Lys and peptide N-termini. Methionine oxidation (+15.9949) was allowed as variable modification. Peptide spectral matches were filtered to a peptide false discovery rate (FDR) of <1% using linear discriminant analysis employing a target-decoy strategy^77,78^. Resulting peptides were further filtered to obtain a 1% protein FDR at the entire dataset level^79^. TMT reporter ion signal-to-noise (S/N) ratios were extracted for quantification^80^.

### Quantitative comparison of proteomes in the clock neuron

Proteins were identified and filtered to retain those with > 2 unique peptides. Quantitative TMT-based ratios were calculated as the fold-change between experimental samples and negative controls. A ratiometric cutoff of log_2_(FC) >=0.5 (equivalent to a ratio >= 1.41) was applied to distinguish bona fide membrane-associated proteins from background contaminants. For Gene Ontology (GO) and Reactome pathway enrichment, identified proteomes were analyzed using the STRING database. The top five enriched terms for cellular compartment and molecular function were plotted based on the lowest false discovery rates (FDR). For the protein-protein interaction (PPI) network (**Figure 3B**), proteins annotated with the "plasma membrane" GO term were extracted. Their interactions were mapped using STRING confidence scores and clustered via the Markov Cluster Algorithm (MCL; inflation value = 1.5).

For the volcano plot (**Figure 3C**), TMT intensities (**Data S1**) for each protein were log_2_ transformed. Proteins with zero or missing values in any channel were excluded. The two negative control (NC) channels from the same time point were averaged (mean of log_2_ values), giving one NC value for dawn and one for dusk. For each protein, the four experimental channels (two replicates each at dawn and dusk) and the two averaged NC values were fitted with a linear model:

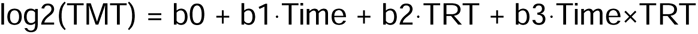

where Time and TRT (treatment) are indicator variables for time of day (Time = 1 for dawn, ZT0; 0 for dusk, ZT11) and labeling condition (TRT = 1 for experimental; 0 for negative control). Each averaged NC value was given a weight of 2 to reflect that it is the mean of two channels. The coefficient b3 is the log_2_ ratio of labeling enrichment (experimental over negative control) at dawn relative to dusk. The null hypothesis that b3 = 0 was tested with a moderated t-test using the limma package in R^81^, with empirical Bayes moderation of the protein-wise variances. Because the NC values were averaged, residual variance was estimated from the experimental replicates. Enriched candidates were additionally checked to be enriched over the negative control at the corresponding time point. Nominal p-values were adjusted for multiple testing by the Benjamini-Hochberg test^82^.

### RNA data processing

Raw count matrices from *Drosophila* clock neuron scRNA-seq datasets (GSE157504; Ma et al., 2021) were obtained from the Gene Expression Omnibus. Corresponding cell annotations and metadata were retrieved from the authors’ public GitHub repository. A Seurat object was constructed using Seurat v5^83^. Metadata were appended to categorize sampling time points into "Early" (ZT2, ZT6, and ZT10) and "Late" (ZT14, ZT18, and ZT22) conditions. Pseudobulk profiles were generated using the ViaFoundry Single-Cell Pseudobulk Workflow (pipeline ID: c73b45866ee645db931a1cd79840c714). Cells were aggregated by time point for light:dark (LD) samples to generate sample-level count vectors. Data were subjected to median ratio normalization (MRN), followed by variance-stabilizing transformation (VST) and log_2_ scaling using DESeq2^84^. Genes annotated as "plasma membrane proteins" were extracted from the top 1,000 transcripts enriched across all clock neurons to facilitate a direct comparison with the proteomic dataset (**Figures. 2D, and 2I**). To account for the inherent temporal lag between transcription and translation, transcriptomic profiles from the "Early" time point were compared against the "dusk" proteome, while "Late" transcriptomic profiles were compared against the "dawn" proteome.

### Locomotor Activity Recording

Male flies (3–5 days old) were individually loaded into glass locomotor tubes containing 4% sucrose and 2% agar. Locomotor activity was recorded using the *Drosophila* Activity Monitoring (DAM) system (TriKinetics, USA), with monitors placed in programmable environmental incubators (Percival Scientific, USA). Activity was recorded in 1-minute bins. For the genetic screen (**Figures 4A, and S3A**), flies were entrained to a 12:12 light:dark cycle at 25°C (LD 25) for 4 days before being released into constant darkness at 25°C (DD 25) for 7 days. Double-plotted actograms, daily activity averages, rhythmicity, and free-running periods were calculated using the FlyToolbox implemented in MATLAB (MathWorks, Inc.)^85^. Period length and significance were determined using the autocorrelation function; flies with an RS value > 1.5 were classified as rhythmic. To test behavioral adaptation to short and long photoperiod conditions, flies were entrained either to a 6:18 (winter-like) or an 18:6 (summer-like) LD cycle at 25°C for 5–7 days. The phase of the morning (M) peak was quantified as the time point of peak average activity between ZT14 and ZT0, while the phase of the evening (E) peak under long photoperiod as peak activity between ZT12 and ZT18.

### Immunohistochemistry

Whole-mount brain immunostaining was performed as previously described^86^. For validation of the biotinylation protocol, flies were entrained under LD 25°C and dissected at ZT8–10. Following fixation in 4% formaldehyde in PBS, brains were washed twice for 10 minutes in PBS, permeabilized in PBS with 0.1% Triton X-100 (PBT) for 30 minutes, and blocked in 10% normal donkey serum (NDS) for 30 minutes. Brains were incubated with primary antibodies (e.g., mouse anti-PDF; 1:500, Developmental Studies Hybridoma Bank) in PBT with 10% NDS overnight at 4°C. After three 10-minute washes in PBT, brains were incubated with secondary antibodies (e.g., Alexa Fluor-488 conjugated anti-mouse, 1:400; Neutravidin-647, 1:400; Invitrogen) in PBT overnight at 4°C in the dark. Slides were washed three times for 5 minutes in PBT and mounted in Vectashield (Vector Laboratories). For Octβ1R and Piezo expression analysis, flies were entrained under LD 25°C for 3 days and dissected at ZT20. Brains were fixed for 20 minutes at room temperature, blocked, and incubated with primary antibodies (rabbit anti-PER^87^, 1:1,000; mouse anti-GFP, 1:500; Invitrogen) followed by Alexa Fluor-conjugated secondary antibodies (Alexa Fluor-488 anti-mouse, 1:500; Alexa Fluor-647 anti-rabbit, 1:1,000; Invitrogen). Images were acquired using a Zeiss LSM 900 laser-scanning confocal microscope (Carl Zeiss).

To quantify s-LNv neural plasticity, control (*w^11^*^18^) and *Piezo^KO^* flies were entrained under LD 25°C and sampled at the indicated time points on the 4^th^ day. PDF-stained axonal arborizations were quantified using the Python-based GUI, *MorphoScope*^88^, which implements the methodology established by^37^. To facilitate temporal comparisons, data were normalized by dividing each time-point value by the peak axonal arborization value.

### CaMPARI Imaging of Ca^2+^ Signals in *Drosophila* Clock Neurons

Flies expressing UAS-CaMPARI2 crossed with *tim-GAL4:16*^89^ in wild-type or *Piezo^KO^* mutant background were used to monitor Ca^2+^ dynamics in clock neurons. Male progenies were entrained either in 12h:12h or 6h:18h light-dark cycles at 25°C for 4–5 days before imaging.

Flies were prepared for imaging by mounting on Petri dishes and stimulating with UV light (395 nm) for 5 minutes. Following photoconversion, their brains were immediately dissected in cold extracellular fly buffer under dim red light. Brain tissues were then imaged using a Zeiss LSM900 confocal microscope.

Analysis involved processing images in ImageJ, with background subtraction to measure the green (F_green_) and red (F_red_) fluorescence intensities in target neurons. The CaMPARI photoconversion efficiency was calculated as the F_red_/F_green_ ratio.

### Statistical Analysis

Statistical analyses of proteomic data were performed in R (version 4.5.1), and behavioral data were plotted and analyzed using GraphPad Prism 11 (GraphPad Software). Data are presented as mean ± SEM unless otherwise stated in the figure legends. Data normality was assessed using the D’Agostino–Pearson omnibus test. For normally distributed data, comparisons between two groups were performed using a two-tailed Student’s t-test, while comparisons among multiple groups were performed using one-way ANOVA followed by Tukey’s multiple-comparisons test. For non-normally distributed data, comparisons between two groups were performed using the Mann–Whitney U test, while comparisons among multiple groups were performed using the Kruskal–Wallis test followed by Dunn’s multiple-comparisons test. Percentage of rhythmic flies was compared between control and mutant flies using a Fisher’s exact test. Statistical significance was defined as P < 0.05, with significance levels indicated as follows: **** P < 0.0001, *** P < 0.001, ** P < 0.01, and * P < 0.05. The statistical tests used for individual experiments are specified in the corresponding figure legends.

## Supporting information

DataS1

DataS2

## Data availability

Original data will be made available upon request.

## Author contribution

Conceptualization, C.C. and P.E.; methodology, C.C., M.G-P., V.K., Q.Y. and P.E.; investigation, C.C., R.C., V.L., L.E.N., Y.X., S.L. and M.G-P..; writing – original draft, C.C.; writing – review & editing, C.C., M.G-P., Q,Y., V.K. and P.E.; funding acquisition, P.E.,V.K., Q.Y.; supervision, P.E., Q.Y. and V.K.

## Acknowledgment

We thank Vinh Phan for technical support. We are grateful to all members of the Emery lab for important discussions and feedback. We also thank the Vienna Drosophila Resource Center and the Bloomington Drosophila Stock Center for providing fly strains. This work was supported by MIRA award 1R35GM145253 from the National Institutes of General Medical Science to PE, and grants from the National Cancer Institute (1K22CA282268), the Worcester Foundation and the American Society for Mass Spectrometry to QY.

## Notes

### Competing Interest Statement

The authors have declared no competing interest.

### Summary of Updates

This revised version of the manuscript includes updated proteomic dataset analysis as well as novel behavioral and neuro-imaging data that demonstrate the importance of newly identified membrane proteins in the control of circadian behavior.

