## Supplementary material for "Membrane proteomics of the *Drosophila* circadian neural network": FigureS1-5TableS1-3

### Figure S1

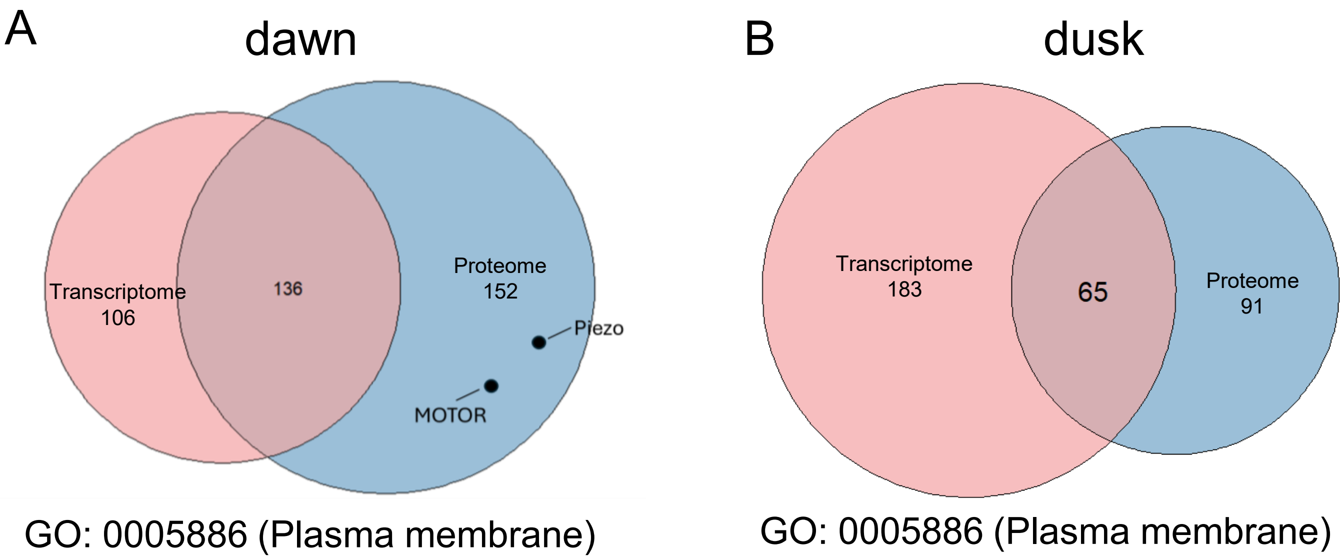


**Supplemental Figure 1. Differences between transcriptome and proteome CMPs in *Drosophila* clock neurons.**

Comparison of the plasma membrane transcriptome (extracted from [1]) versus the proteome of circadian neurons. Note the differences between mRNA and protein enrichment for a significant fraction of the libraries. Related to **Figure 3**.

Figure S2

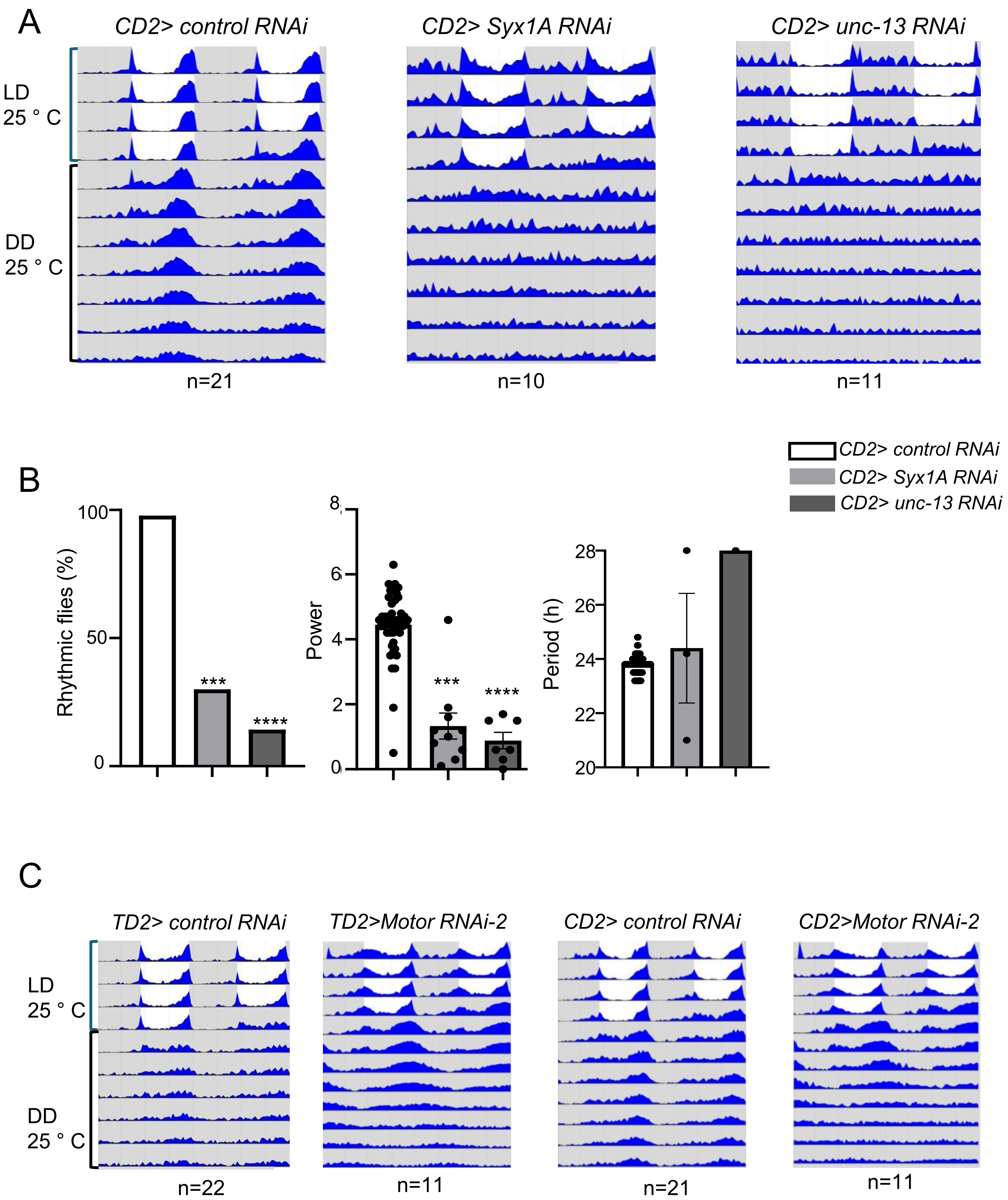


**Supplemental Figure 2. Knockdown of cell membrane proteins compromise behavioral rhythms under constant darkness (DD)**

(A) Knockdown of *Syx1A* and *unc-13* via *Clk-GAL4* (CD2) severely decrease rhythmicity in DD. Representative double plotted actogram displaying behavior under LD and DD are shown for control (*mCherry RNAi*), *Syx1A* RNAi, and *unc-13* RNAi flies. (B) Quantification of behavior for the genotypes shown in (A). For period length, only rhythmic flies were included. The percentage of rhythmic flies was compared with controls using a chi-square test ***p < 0.001,****p < 0.0001. Rhythmic power and period length were evaluated using a Kruskal-Wallis test and Dunn's post-hoc test, ***p < 0.001,****p < 0.0001. (C) Knockdown of *Motor* (*CG3638*) with a second RNAi transgene also leads to a significantly lengthened period in DD. Related to **Figure 4**.

Figure S3

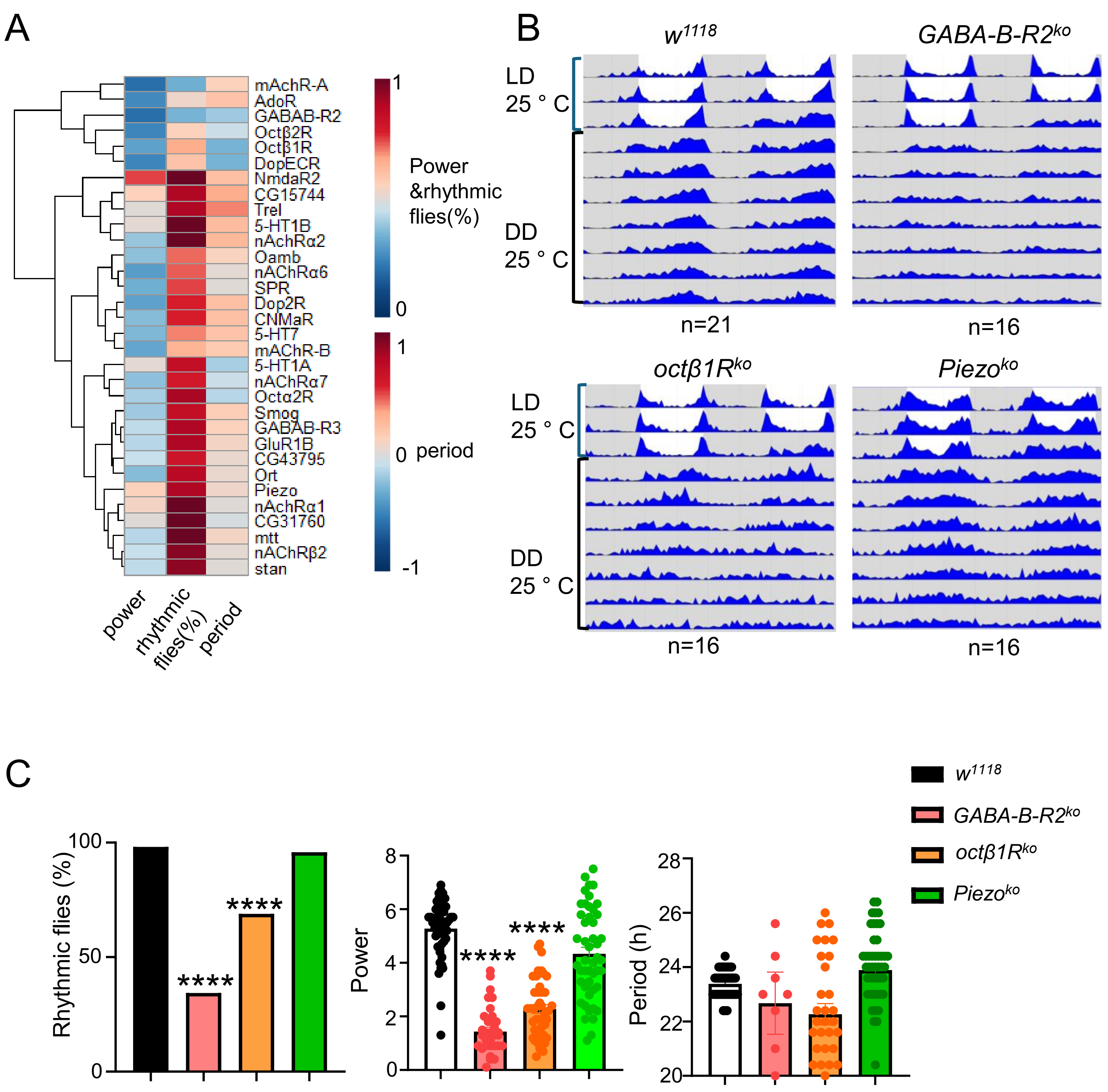


**Supplemental Figure 3. A circadian behavior proteome-guided genetic screen using knockout mutants .**

(A) Heatmap summary of normalized rhythmicity percentage, power, and period length in DD using null mutation of candidate genes. Data are normalized as described in **Figure 4A**. Each candidate was tested with n >=10 flies. (B) Knockout of *GABA-B-R2* results in arrhythmicity. Double plotted actograms in LD and DD are shown for control ($w^{1118}$), *GABA-B-R2*$^{ko}$, *Octβ1R*$^{ko}$, and *Piezo*$^{KO}$. (C) Quantification of behavior for the genotypes in (B). For period length, only rhythmic flies were included. For percentage of rhythmic flies, a Fisher's exact test was used to compare each mutant to control flies, ***$p < 0.001$,****$p < 0.0001$, For rhythmic power and period length, Kruskal-Wallis test and Dunn's post-hoc test were used, ***$p < 0.001$,****$p < 0.0001$. Related to **Figures 4, 5 and 6.**

Figure S4

A

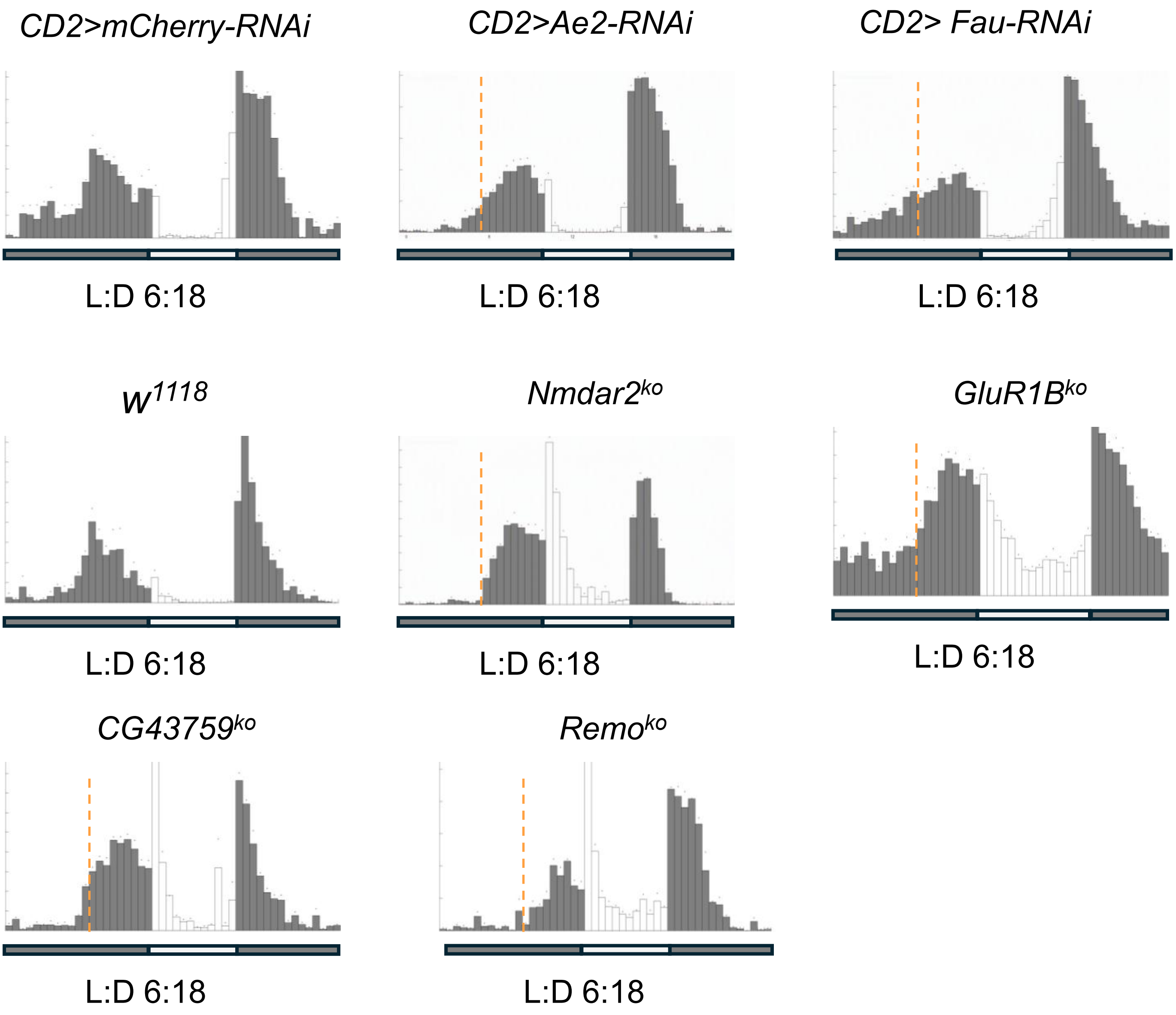


**Supplemental Figure 4. Genetic screen for CMP candidates under short photoperiod.** Representative daily locomotor activity averages of control and mutant flies (n>=8) under short photoperiod. Orange dash lines indicate the position of the morning activity peak in control flies. Related to **Figure 6.**

Figure S5

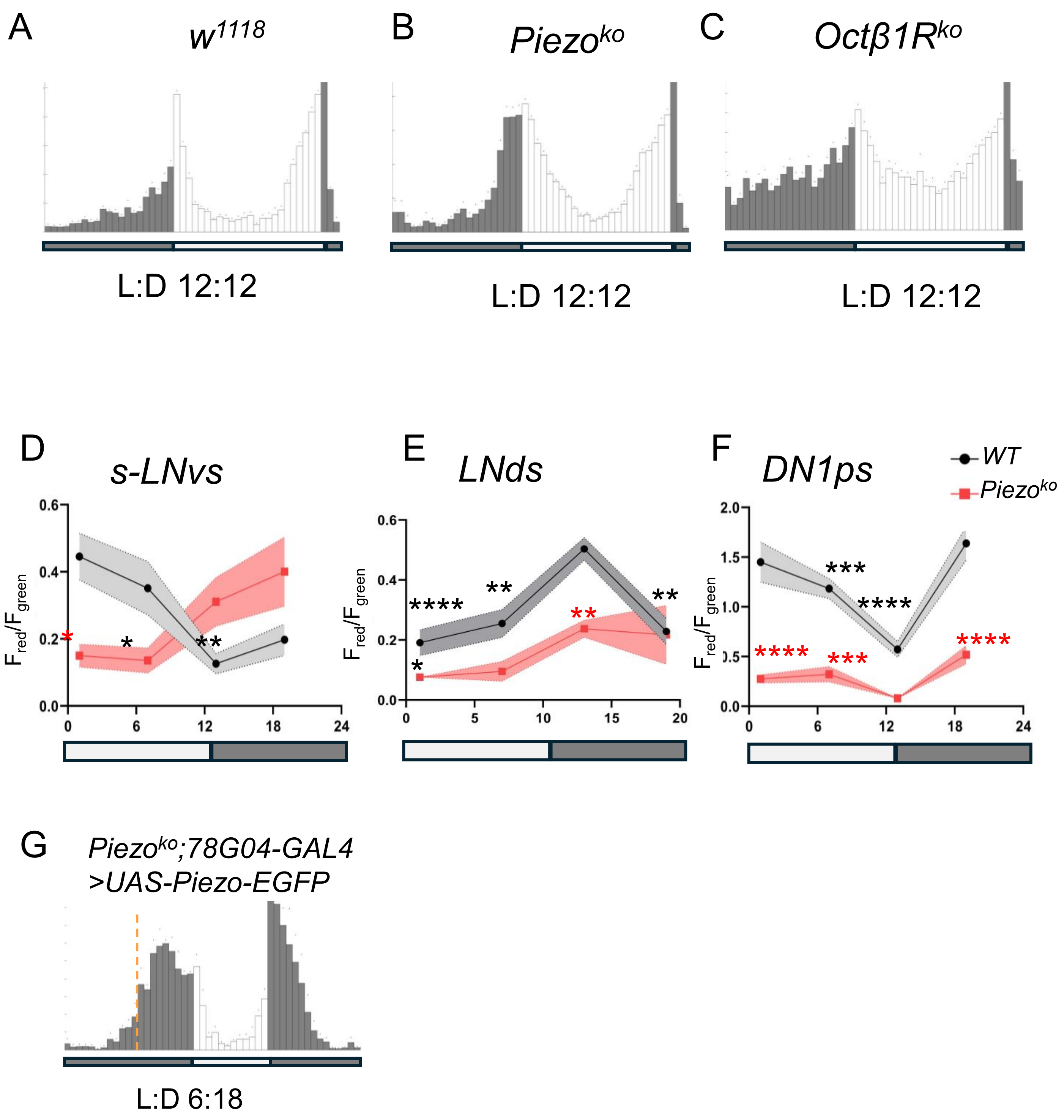


**Supplemental Figure 5. Octβ1R and Piezo are not required for behavioral adaptation to L:D 12:12.**

(A-C) Representative locomotor activity of *w*$^{1118}$(n=11), *Octβ1R*$^{KO}$ (n=11) and *Piezo*$^{KO}$ (n=11) under normal photoperiod. (D-F) Same as in **Figure 7D-F**, but quantification of $Ca^{2+}$ signals using CaMPARI2 in subsets of clock neurons in WT and *Piezo*$^{KO}$ flies under normal photoperiod (L:D, 12:12), error bars, SEM. ****P < 0.001, ***P < 0.001, **P < 0.01, and * P < 0.05, two-way ANOVA with Tukey's post hoc test. Red asterisks indicate significant differences between WT and mutant flies, whereas black asterisks represent significant differences relative to the peak value within the same genotype (G) Restoration of wild-type *Piezo* in l-LNvs (with the *78G04-GAL4* driver) failed to rescue the mutant phenotype. Dash line (orange) indicates morning activity peak of control flies (*w*$^{1118}$) under short photoperiod condition. Related to **Figures 5, 6 and 7.**

**Table S1. Membrane proteins previously associated with Drosophila circadian rhythms identified as highly enriched in clock neurons**

| Gene | Molecular Function | Express in clock neuron | Function in clock/sleep | References |
|---|---|---|---|---|
| *Gγ30A* | G protein complex subunit | yes | | [2] |
| *brp* | Structure for synapse | | yes | [3] |
| *cac* | voltage-gated $Ca^{2+}$channel subunit | | yes | [4] |
| *dysc* | Regulator of $Ca^{2+}$-activated $K^{+}$ channel | | yes | [5] |
| *Dgk* | Downstream of GPCR | yes | | [2] |
| *GABA-B-R1* | GPCR of GABA | yes | | [6,7] |
| *Octβ2R* | GPCR of octopamine | yes | | [7,8] |
| *Ih* | voltage-gated ion channel | | yes | [9] |
| *SK* | $Ca^{2+}$-activated $K^{+}$ channel | yes | yes | [10] |
| *Shaw* | voltage-gated $K^{+}$ channel | yes | yes | [11,12] |
| *Dlg1* | Synaptic guanylate kinase | yes | | [2] |
| *Oct-TyrR* | Receptor of Tyramine | yes | | [8] |
| *mAChR-A* | Receptor for muscarinic acetylcholine | yes | | [13] |
| *slo* | Subunit of calcium-activated potassium channel | | yes | [14] |
| *rut* | Downstream of GPCR | | yes | [15] |
| *nAChRα1* | nicotinic acetylcholine receptor | yes | | [6,7] |
| *mtt* | GPCR | yes | | [6] |
| *Inx2* | Gap junction | yes | yes | [16] |
| *Ndae1* | $Na^{+}$-driven anion exchanger | yes | | [2] |
| *GABA-B-R2* | GABA receptor | yes | | [6,7] |
| *nrv1* | Na+, K+ ATPase ion pump | yes | | [2] |
| *na* | Sodium leak channel | yes | yes | [17,18] |
| *Alk* | Ras/ERK and PI3K signaling pathways | | yes | [19] |
| *Ca-α1T* | voltage gated calcium channel subunit | yes | yes | [20] |
| *Dop1R1* | GPCR of Dopamine | yes | | [13,21] |

**Table S2. RNAi screen under DD of top enriched membrane proteins**

| Gene | Stock number | Rhythmic flies% | Power ±SEM | Period (h) ±SEM |
|---|---|---|---|---|
| *mCherry-RNAi* | BDSC#35785 | 100 | 4.0±0.3 | 23.7± 0.1 |
| *nAchRβ1* | BDSC#31883 | 100 | 4.2±0.3 | 23.7± 0.1 |
| *Ndae1* | BDSC#62177 | 55.6 ** | 2.0± 0.5 | 24.0 ±0.3 |
| *CG1909* | BDSC#28024 | 80.0 | 3.5± 0.5 | 23.8± 0.1 |
| *5-HT2B* | BDSC#25874 | 87.5 | 3.8± 0.6 | 24.0± 0.2 |
| *Con* | BDSC#28967 | 80.0 | 2.7± 0.4 | 23.8 ±0.3 |
| *nAchRα4* | BDSC#31985 | 90.9 | 3.6± 0.37 | 23.9± 0.1 |
| *slo* | BDSC#26247 | 70.0* | 2.6± 0.4 | 23.8± 0.2 |
| *ogre(inx1)* | BDSC#27283 | 60.0** | 1.6± 0.2** | 23.4± 0.8 |
| *unc-13* | BDSC#29548 | 27.3**** | 1.3± 0.2*** | 21.8 ±0.7 |
| *ELK* | BDSC#25821 | 55.6** | 2.1 ±0.6 | 23.8± 0.1 |
| *Gluclα* | BDSC#53356 | 90.0 | 2.7 ±0.3 | 23.4± 0.1 |
| *Shawl* | BDSC#25819 | 70.0* | 2.7± 0.5 | 23.8 ±0.1 |
| *tutl* | BDSC#54850 | 100.0 | 4.8± 0.3 | 23.8± 0.1 |
| *Nhe2* | BDSC#51491 | 100.0 | 3.7 ±0.4 | 23.9± 0.2 |
| *TM4SF* | BDSC#39048 | 80.0 | 2.8± 0.5 | 23.5± 0.2 |
| *bdl* | BDSC#31974 | 50.0** | 1.0 ±0.6* | 24.2 ±0.2 |
| *αPKC* | BDSC#25946 | 100.0 | 4.4± 0.3 | 23.9 ±0.2 |
| *PsGEF* | BDSC#33433 | 100.0 | 3.3± 0.3 | 24.1± 0.2 |
| *Inx2* | BDSC#29306 | 77.8* | 2.0± 0.2* | 23.7 ±0.2 |
| *PMCA* | BDSC#31572 | 100.0 | 4.8± 0.3 | 24.6 ±0.2* |
| *Vha100-1* | BDSC#57860 | 25.0**** | 1.1± 0.3** | 24.8± 3.0 |
| *Syx1A* | BDSC#25811 | 30.0**** | 1.3 ±0.4** | 24.4± 2.0 |
| *fau* | BDSC#60351 | 90.9 | 4.1± 0.3 | 24.3± 0.1 |
| *Hk* | BDSC#28330 | 100.0 | 3.6 ±0.3 | 23.9± 0.1 |
| *Dgk* | BDSC#29459 | 77.8* | 2.3 ±0.4 | 24.3 ±0.2 |
| *NaCp60E* | BDSC#26012 | 100.0 | 3.2 ±0.3 | 23.9 ±0.1 |
| *patj* | BDSC#26282 | 90.0 | 3.7 ±0.3 | 24.2 ±0.1 |
| *metro* | BDSC#35810 | 88.9 | 3.2 ±0.4 | 23.9± 0.1 |
| *cno* | BDSC#38194 | 100.0 | 4.8 ±0.2 | 24.2 ±0.1 |
| *elm* | BDSC#40843 | 100.0 | 3.6 ±0.3 | 24.0± 0.2 |
| *α-Cat* | BDSC#33430 | 100.0 | 4.7± 0.3 | 24.1 ±0.1 |
| *Flo1* | BDSC#36649 | 100.0 | 4.8± 0.2 | 23.9± 0.1 |
| *kcc* | BDSC#34584 | 100.0 | 4.3± 0.7 | 24.2± 0.1 |
| *hzg* | BDSC#31746 | 100.0 | 4.6± 0.3 | 23.8 ±0.1 |
| *VhaAC45RP* | BDSC#64549 | 100.0 | 4.2± 0.5 | 23.4± 0.2 |
| *Ih* | BDSC#29574 | 100.0 | 3.4± 0.3 | 23.8 ±0.1 |
| *Nrv2* | BDSC#28666 | 87.5 | 3.6 ±0.5 | 24.0± 0.1 |
| *Sh* | BDSC#53347 | 100.0 | 4.3 ±0.2 | 24.8± 0.1*** |
| *Ca-α1D* | BDSC#25830 | 100.0 | 3.7± 0.3 | 23.7 ±0.1 |
| *Ca-β* | BDSC#29575 | 90.9 | 3.6± 0.5 | 23.6 ±0.2 |
| *Prip* | BDSC#50695 | 100.0 | 4.1 ±0.4 | 23.9 ±0.2 |
| *ShakB* | BDSC#27291 | 100.0 | 3.9 ±0.3 | 23.6 ±0.1 |
| *StmA* | BDSC#39062 | 81.8 | 2.5 ±0.4 | 23.8± 0.2 |
| *Irk2* | BDSC#81688 | 100.0 | 3.8 ±0.2 | 23.7± 0.1 |
| *P120ctn* | BDSC#51729 | 100.0 | 4.8 ±0.2 | 23.7± 0.1 |
| *Shal* | BDSC#31879 | 100.0 | 4.4± 0.2 | 24.0± 0.1 |
| *Dap160* | BDSC#25879 | 77.8* | 2.7± 0.6 | 24.2 ±0.2 |
| *Motor* | VDRC#102444 | 90.9 | 3.3± 0.4 | 25.8 ±0.2**** |
| *CG42748* | VDRC#107426 | 55.6 | 2.9 ±0.6 | 24.0± 0.2 |
| *VhaM9.7-a* | VDRC#104315 | 100.0 | 2.6± 0.3 | 24.2 ±0.2 |
| *tty* | VDRC#110431 | 90.0 | 3.7 ±0.5 | 24.6± 0.2* |
| *Ac76E* | VDRC#106232 | 100.0 | 3.5± 0.6 | 24.6 ±0.4 |
| *CG42258* | VDRC#109934 | 80.0 | 3.5± 0.6 | 24.5 ±0.2 |
| promL | VDRC#102612 | 90.9 | 4.4± 0.4 | 24.3 ±0.1 |
| Teh1 | VDRC#102816 | 100.0 | 4.3 ±0.2 | 24.4 ±0.1* |

**Table S2. RNAi screen under DD of top enriched membrane proteins-continued**

| Gene | Stock number | Rhythmic flies% | Power ±SEM | Period (h) ±SEM |
|---|---|---|---|---|
| *Zasp52* | BDSC#58198 | 100.0 | 4.1 ±0.2 | 24.4± 0.1 |
| *Akap200* | BDSC#28532 | 90.9 | 3.2 ±0.3 | 24.2± 0.2 |
| *mbc* | BDSC#33722 | 87.5 | 3.6 ±0.5 | 23.5 ±0.1 |
| *CG43066* | BDSC#28069 | 100.0 | 3.5 ±0.3 | 24.2± 0.1 |
| *CG8245* | BDSC#29426 | 10.0**** | 1.1 ±0.1*** | 27.8±0.0 |
| *NMDAR1* | BDSC#25941 | 100.0 | 3.3 ±0.3 | 24.4 ±0.2 |
| *Rdl* | BDSC#41101 | 100.0 | 4.4± 0.2 | 23.9± 0.1 |
| *Lar* | BDSC#34965 | 66.7* | 1.5± 0.2* | 24.1 ±2.0 |
| *Tomosyn* | BDSC#31980 | 100.0 | 4.4± 0.3 | 24.3 ±0.2 |
| *PHcl-1* | BDSC#38909 | 100.0 | 4.6± 0.2 | 24.2 ±0.2 |
| *Rh50* | BDSC#50666 | 100.0 | 4.0± 0.3 | 24.2 ±0.1 |
| *Dlg1* | BDSC#25780 | 100.0 | 4.0± 0.6 | 24.0 ±0.1 |
| *Itpr* | BDSC#25937 | 100.0 | 3.6± 0.4 | 23.4 ±0.1 |
| *Syt7* | BDSC#27279 | 100.0 | 3.7 ±0.4 | 24.1 ±0.1 |
| *Ae2* | BDSC#51827 | 100.0 | 4.2 ±0.1 | 24.3± 0.1 |
| *bark* | BDSC#67014 | 100.0 | 4.2 ±0.2 | 24.3± 0.1 |
| *RSG7* | BDSC#28574 | 87.5 | 3.4 ±0.4 | 24.0 ±0.2 |
| *Atpα* | BDSC#32913 | 100.0 | 3.5 ±0.4 | 23.6 ±0.1 |
| *SK* | BDSC#27238 | 100.0 | 3.7 ±0.4 | 23.7 ±0.2 |
| *Eglp4* | BDSC#67959 | 100.0 | 4.3± 0.2 | 23.9 ±0.1 |
| *SLO2* | BDSC#32034 | 100.0 | 3.8 ±0.3 | 23.8 ±0.1 |

All genotypes include thetim-GAL4driver and the UAS-dcr2 transgene. Percentage of rhythmic flies, period and power of rhythmicity were determined as described in the Method section. UAS-mCherry RNAi flies served as the control group. A Fisher's exact test was used to determine the statistical significance of differences in percentage of rhythmic flies, while a Kruskal–Wallis test followed by Dunn's multiple-comparisons test was used to determine the significance of changes in rhythmic power and period length. ****: $p <0.0001$, ***: $p < 0.001$, **: $p < 0.01$, *: $p < 0.05$.

**Table S3. Gene knockout screen under DD of top enriched membrane proteins**

| **gene** | **stock number** | **rhythmic flies%** | **Power** ± SEM | **Period (h)** ±SEM |
|---|---|---|---|---|
| *w*$^{1118}$ | Lab stock | 100 | 5.3±0.1 | 23.7±0.6 |
| *GABABR2* | BDSC#84504 | 66.7$^{****}$ | 1.4±0.2$^{****}$ | 23.3±0.6 |
| *GABABR3* | BDSC#84505 | 90.0 | 3.2±0.3$^{***}$ | 23.9±0.2 |
| *5-HT1A* | BDSC#84704 | 88.9 | 3.7±0.4 | 23.4±0.1 |
| *5-HT1B* | BDSC#84443 | 100.0 | 3.7±0.4 | 24.1±0.1 |
| *5-HT7* | BDSC#84446 | 77.8$^{*}$ | 2.6±0.5$^{***}$ | 24.0±0.3 |
| *nAChRα7* | BDSC#84543 | 80.0$^{*}$ | 2.7±0.4$^{**}$ | 23.6±0.2 |
| *nAchRα1* | BDSC#84539 | 100.0 | 3.9±0.5 | 23.7±0.1 |
| *nAchRα2* | BDSC#84540 | 93.3 | 2.7±0.3$^{****}$ | 23.9±0.1 |
| *nAChRβ2* | BDSC#84545 | 97.4 | 3.7±0.2$^{**}$ | 23.6±0.1 |
| *nAChRα6* | BDSC#84542 | 72.2$^{***}$ | 2.7±0.2$^{****}$ | 23.7±0.2 |
| *mAchRA* | BDSC#84519 | 33.3$^{****}$ | 1.5±0.3$^{****}$ | 24.0±2.0 |
| *mAChRB* | BDSC#84520 | 63.2$^{****}$ | 2.3±0.4$^{****}$ | 24.0±0.2 |
| *Octα2R* | BDSC#84472 | 55.6$^{***}$ | 3.0±0.3$^{**}$ | 23.8±0.1 |
| *Octβ1R* | BDSC#84554 | 70.5$^{****}$ | 2.3±0.2$^{****}$ | 23.5±0.1 |
| *Octβ2R* | BDSC#84545 | 55.6$^{***}$ | 1.8±0.4$^{****}$ | 23.1±0.2 |
| *stan* | BDSC#84577 | 94.7 | 3.2±0.2$^{**}$ | 23.7±0.1 |
| *CG31760* | BDSC#84474 | 100.0 | 3.6±0.4 | 23.6±0.2 |
| *Dop2R* | BDSC#84495 | 77.8$^{*}$ | 2.2±0.7$^{***}$ | 24.0±0.1 |
| *DopECR* | BDSC#84496 | 59.1$^{****}$ | 1.8±0.2$^{****}$ | 23.2±0.2 |
| *Remo* | BDSC#97780 | 90.0 | 4.0±0.3 | 24.2±0.1$^{**}$ |
| *Smog* | BDSC#97853 | 84.2 | 2.9±0.3$^{***}$ | 23.9±0.3 |
| *Trel* | BDSC#84582 | 90.0 | 3.6±0.5 | 23.3±1.2 |
| *AdoR* | BDSC#84447 | 51.7$^{****}$ | 1.8±0.2$^{****}$ | 24.2±0.3 |
| *CG43795* | BDSC#84480 | 81.8$^{*}$ | 3.3±0.4$^{*}$ | 23.8±0.1 |
| *Ort* | BDSC#84559 | 87.5 | 2.7±0.4$^{**}$ | 23.8±0.3 |
| *SPR* | BDSC#84692 | 75.0$^{**}$ | 2.5±0.2$^{****}$ | 23.7±0.3 |
| *NmdaR2* | BDSC#84548 | 100.0 | 5.5±0.2 | 24.1±0.1 |
| *GluR1B* | BDSC#84507 | 89.5 | 3.1±0.3$^{***}$ | 23.8±0.1 |
| *CNMaR* | BDSC#84486 | 77.8$^{*}$ | 2.7±0.5$^{**}$ | 24.1±0.3 |
| *Oamb* | BDSC#84555 | 71.4$^{*}$ | 2.7±0.7$^{*}$ | 23.8±0.2 |
| *mtt* | BDSC#84538 | 93.8 | 2.9±0.3$^{***}$ | 23.8±0.3 |
| *Piezo* | BDSC#58770 | 95.8 | 4.3±0.2 | 24.0±0.1 |

Percentage of rhythmic flies, period and power of rhythmicity were determined as described in the Method section. w1118 flies served as the positive control. A Fisher's exact test was used to determine the statistical significance of differences in percentage of rhythmic flies, while a Kruskal–Wallis test followed by Dunn's multiple-comparisons test was used to determine the significance of changes in rhythmic power and period length. ****: $p < 0.0001$, ***: $p < 0.001$, **: $p < 0.01$, *: $p < 0.05$.
